# LRRK2 regulates AP2M1 phosphorylation cycles to mediate endocytosis and dopaminergic neurodegeneration

**DOI:** 10.1101/2020.05.27.119388

**Authors:** Qinfang Liu, Judith Bautista-Gomez, Daniel A. Higgins, Jianzhong Yu, Yulan Xiong

**Author notes:** To whom correspondence should be addressed: Yulan Xiong, or Jianzhong Yu, Department of Anatomy & Physiology, Kansas State University College of Veterinary Medicine, Manhattan, KS 66506, USA.

## Abstract

Recent genetic evidence revealed endocytic pathway plays a major role in Parkinson’s disease (PD). However, the molecular mechanism is poorly understood. Here we report that LRRK2, the most genetic cause of PD, binds to and phosphorylates AP2M1, the core component of endocytosis recently implicated in PD risk. Both knockout and overexpression of LRRK2 cause abnormal AP2M1 phosphorylation cycle and in turn endocytic defects. Mechanistically, knockout of LRRK2 decreases AP2M1 phosphorylation required for the initial clathrin coated vesicle (CCV) formation while LRRK2 overexpression inhibits AP2M1 uncoating for entering into a new cycle of CCV formation. Our study also uncovered a novel tissue-specific regulation of AP2M1 phosphorylation by LRRK2. Further, we found LRRK2 phosphorylation on AP2M1 mediates LRRK2-induced neuronal toxicity both *in vitro* and *in vivo*. Importantly, AP2M1 phosphorylation levels are elevated in PD patient fibroblasts. Together, our study provides a direct mechanistic link between LRRK2, AP2 and endocytosis in PD pathogenesis.

## Introduction

Advances in genetics and multiple model systems have revealed an important role of intracellular trafficking defects in Parkinson’s disease (PD)(1–3). Importantly, a recent systematic analysis of pathway-specific genetic risk factors revealed that endocytic membrane-trafficking pathway plays a major role in the risk of PD(2). Moreover, leucine rich repeat kinase 2 (LRRK2), the mutations in which are the most genetic causes of both familial and sporadic PD(4, 5), has been implicated in the vesicle trafficking including endocytosis(1, 3, 6, 7). Specific expression of the most prevalent LRRK2 mutation G2019S in dopaminergic neurons in mouse brain induces a robust reduction in the number of synaptic vesicles and an accumulation of clathrin coated vesicles (CCVs) at synapses, suggesting a clathrin-mediated endocytic (CME) trafficking defect *in vivo*(8). Despite these advances, our understanding of how LRRK2 regulates endocytosis and how the endocytic defects caused by LRRK2 mutations contribute to dopaminergic neurodegeneration remains fragmented.

Neuronal activity is highly reliant on efficient CME to retrieve and maintain synaptic vesicle proteins on the plasma membrane. Therefore, proper regulation of key proteins related to this process is critical for normal neuronal activity. The adaptor protein complex 2 (AP2) and clathrin constitute the major coat constituents in CME(9). AP2 is a heterotetrameric complex and is required for binding the clathrin coat to the membrane and recruiting cargoes into the pit. It consists of two large subunits (α or A1/2, and β2 or B1), a medium subunit (μ2 or M1), and a small subunit (σ2 or S1)(10–12). The large subunits bind to clathrin and recruit other accessory proteins in plasma membrane targeting. The M1 subunit interacts with cargo molecules including transferrin (Tfn) receptor. And the S1 subunit appears to stabilize the AP2 complex(12, 13). Proper phosphorylation-dephosphorylation cycle of the AP2 complex has been implicated in synaptic vesicle recycling and is required for efficient endocytosis(13). Phosphorylation of AP2M1 is critical for its membrane association and initial clathrin coated vesicle (CCV) formation. After CCV scission, AP2M1 dephosphorylation promotes its uncoating from CCVs, a critical process required for a new cycle of CCV formation. Interestingly, AP2M1 has been recently indicated to be relevant for PD risk(2). However, little is known about the underlying mechanisms and how it contributes to dopaminergic neurodegeneration.

Here we found that LRRK2 specifically phosphorylates AP2M1 at T156. Our study demonstrated that ectopic LRRK2 kinase activity inhibits AP2M1 uncoating after CCV scission, leading to the inhibition of subsequent new cycle of CCV formation and endocytosis defects. Interestingly, we found that knockout of LRRK2 also causes endocytosis defects, accompanied with decreased AP2M1 phosphorylation, suggesting basal level LRRK2 kinase activity is required for AP2M1 phosphorylation induced initial CCV formation. Strikingly, we found that the effect of LRRK2 on AP2M1 phosphorylation is dramatically observed in mouse brain, where LRRK2 is abundantly expressed, suggesting a tissue-specific role of LRRK2 on AP2M1 phosphorylation. We further showed that phosphorylation of AP2M1 by LRRK2 mediates LRRK2-induced neuronal toxicity both *in vitro* in neuronal cultures and *in vivo* in *Drosophila*. Importantly, AP2M1 phosphorylation levels are increased in patient fibroblasts of both LRRK2-associated PD and sporadic PD. Taken together, our study revealed a novel mechanistic link between LRRK2, AP2M1 and endocytosis in PD pathogenesis.

## Results

### AP2M1 interacts with LRRK2

In an effort to investigate how LRRK2 mutations mediate cellular trafficking, we performed an unbiased genome-wide genetic screen modified from our previous genetic screen(14), using LRRK2 fragment including the central GTPase, COR, kinase domains (GTP-COR-Kin), and identified yeast deletion mutants that suppress or enhance LRRK2-induced toxicity with a focus on cellular trafficking. Deletion mutants of yeast AP2 four subunits (*apl3*Δ, *apl1*Δ, *apm4*Δ, *aps2*Δ) were identified to rescue LRRK2-induced toxicity and were further confirmed by cell viability assays (Figure 1A). Based on the results from yeast screen, we examined the physical interactions between LRRK2 and the four subunits of human AP2 (A2, B1, M1, S1) by co-immunoprecipitation (co-IP) using overexpressed MYC-LRRK2 and respective HA-AP2 subunits in HEK 293T cells. LRRK2 preferentially pulled down AP2A2 and AP2M1 in cells (Figure 1B). To further determine if LRRK2 directly interacts with AP2A2 and AP2M1 in *in vitro* system, we purified GST-tagged recombinant proteins GST-AP2A2 and GST-AP2M1, and incubated them with purified MYC-LRRK2. IP with anti-MYC strongly coimmunoprecipitates GST-AP2M1 but not GST-AP2A2, nor the GST protein as a negative control (Figure 1C). Thus, we concluded that LRRK2 preferentially interacts with AP2 subunit AP2M1. We then tested if LRRK2 familial mutants differentially interact with AP2M1. We found no significant difference in binding between AP2M1 and wild type (WT) LRRK2 or the familial mutations R1441C (RC), R1441G (RG), G2019S (GS), or I2020T (IT) LRRK2 (Supplemental Figure 1A, B). To examine whether LRRK2 interacts with AP2M1 *in vivo* in mouse brain, we conducted co-IP in whole brain lysates from LRRK2 WT and knockout (KO) mice. Endogenous LRRK2 pulled down endogenous AP2M1 in LRRK2 WT but not KO mice (Figure 1D). To determine the domain of AP2M1 that interacts with LRRK2, we generated AP2M1 deletion mutations at its four structural domains according to SCOP (Structural Classification of Proteins): D1, D2, D3, D4 and examined their interactions with LRRK2. D2 domain but not other three domains of AP2M1 interacts with LRRK2 (Figure 1E). Moreover, the binding domain on LRRK2 that interacts with AP2M1 was determined using our available deletion mutation series spanning the entire LRRK2 proteins with different functional domains(15). We found that AP2M1 predominantly binds to the GTPase domain of LRRK2 (Figure 1F). We further tested whether LRRK2 GTP binding mutants interact with AP2M1. Interestingly, we found AP2M1 does not bind to LRRK2 GTP binding deficient mutant T1348N, and has a slightly increased binding with LRRK2 GTP binding active mutant R1398L, suggesting that LRRK2 GTP binding activity affects the interaction between AP2M1 and LRRK2 (Supplemental Figure 1C). Taken together, LRRK2 preferentially interacts with AP2M1 of AP2 complex.

**Figure 1.**
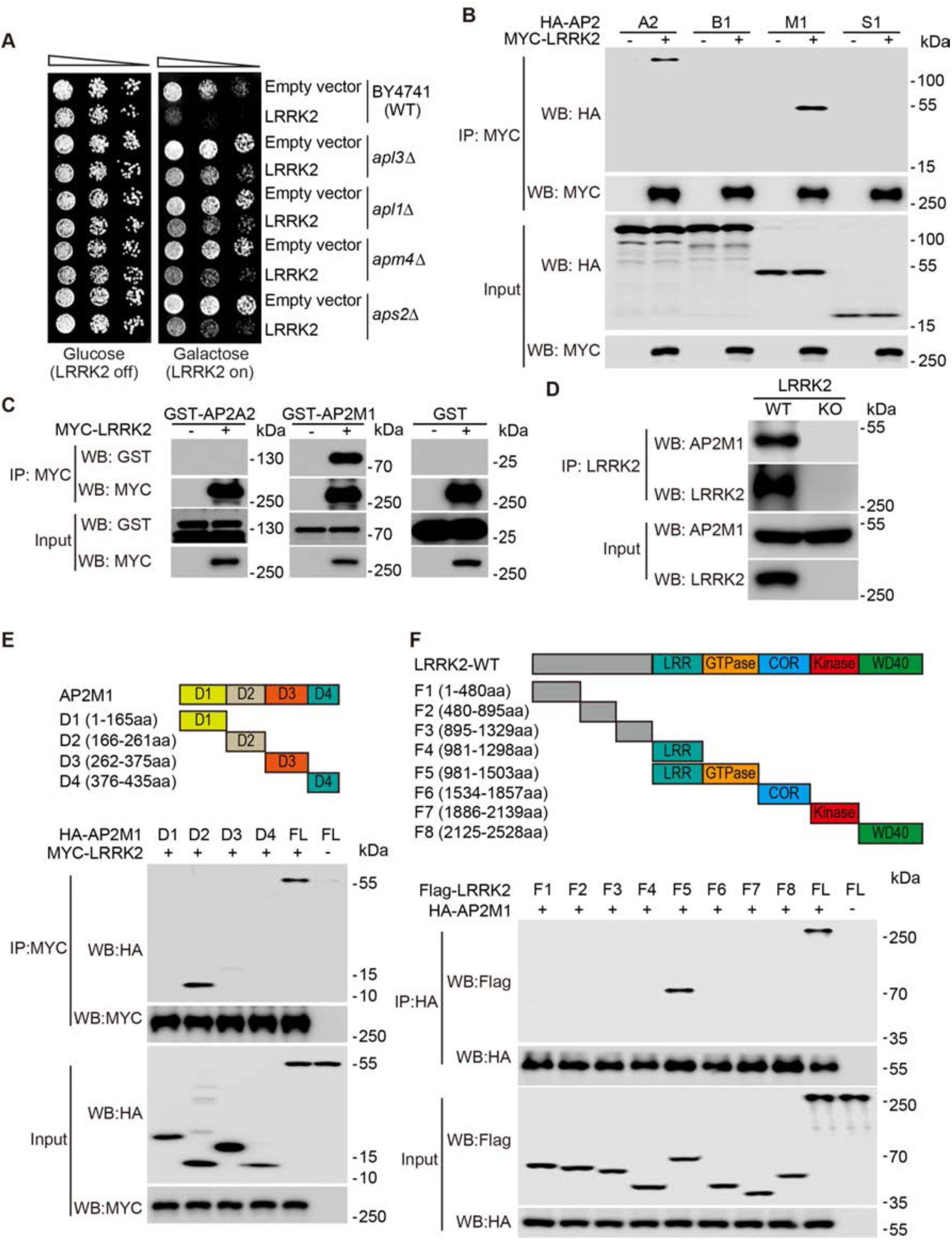
AP2M1 interacts with LRRK2. (**A**) Yeast gene deletion strains of AP2 four subunits strains (*apl3*Δ, *apl1*Δ, *apm4*Δ, *aps2*Δ) markedly rescue LRRK2-induced toxicity compared to the WT strain. Yeast WT strain BY4741 and gene deletion strains *apl3*Δ, *apl1*Δ, *apm4*Δ, *aps2*Δ cells were transformed with galactose-inducible expression constructs containing the central GTP-COR-Kin fragment of WT LRRK2 or empty vector as a control. Cells were spotted onto media containing glucose (LRRK2 Off, repressed, left panel) or galactose (LRRK2 On, induced, right panel) and incubated at 30°C for 2–3 days. Shown are five-fold serial dilutions of yeast cells. From left to right, as indicated by graded open box, starting with equal numbers of cells. (**B**) AP2 subunits A2 and M1 interact with LRRK2 in HEK 293T cells. Cell lysates from HEK 293T cells co-transfected with MYC-LRRK2 and HA-AP2 four subunits (A2, B1, M1, S1) respectively were subjected to coimmunoprecipitation (Co-IP) with anti-MYC followed by anti-HA immunoblotting or by anti-MYC to show an equivalent amount of input. (**C**) Recombinant GST-AP2M1 directly interacts with LRRK2 *in vitro*. Purified GST-tagged recombinant proteins GST-AP2A2 or GST-AP2M1 were incubated with purified MYC-LRRK2 and subjected to Co-IP with anti-MYC followed by anti-GST immunoblotting. GST protein was as a negative control. (**D**) AP2M1 interacts with LRRK2 *in vivo*. Whole brain lysates prepared form wildtype (WT) and LRRK2 knockout (KO) mice were subjected to Co-IP with anti-LRRK2 (Neuromab) followed by anti-LRRK2 and anti-AP2M1 immunoblotting. (**E**) LRRK2 interacts with the D2 domain of AP2M1. Cell lysates from HEK 293T cells co-transfected with MYC-LRRK2 and HA-AP2M1 D1, D2, D3, D4 or full-length (FL) were subjected to Co-IP with anti-MYC followed by anti-HA immunoblotting or by anti-MYC to show an equivalent amount of input. A schematic representation of AP2M1 D1, D2, D3, D4 domains is shown. (**F**) AP2M1 interacts with the GTPase domain of LRRK2. Cell lysates from HEK 293T cells co-transfected with HA-AP2M1 and flag-tagged LRRK2 fragments were subjected to Co-IP with anti-HA followed by anti-flag immunoblotting or by anti-HA to show an equivalent amount of input. A schematic representation of the different LRRK2 fragments used is shown.

### LRRK2 phosphorylates AP2M1 *in vitro* and *in vivo*

LRRK2 mutations are the most common genetic causes of both familial and sporadic PD and LRRK2 kinase activity is highly relevant to PD pathogenesis. Interestingly, proper phosphorylation of AP2M1 is critical for its function in endocytosis. Based on our observed strong interaction between LRRK2 and AP2M1, we asked whether LRRK2 phosphorylates AP2M1. To answer this question, we first performed an *in vitro* kinase assay with [^32^Pγ]-ATP using purified recombinant proteins of LRRK2 and AP2M1. We found that kinase active G2019S (GS) LRRK2 but not kinase dead D1994A (DA) LRRK2 mutant phosphorylates AP2M1 *in vitro* (Figure 2A). A known LRRK2 kinase substrate Rab8a was included as a positive control. To determine the phosphorylation sites, we further subjected the phosphorylated AP2M1 by LRRK2 to mass spectrometry (MS) analysis and identified threonine 156 (T156) of AP2M1 as the major residue phosphorylated by LRRK2 (Supplemental Figure 2A, B). Accordingly, we generated a phosphor-deficient AP2M1-T156A (TA) mutant. Using a phosphor-specific antibody against AP2M1-T156, we further confirmed that WT LRRK2 phosphorylates AP2M1 WT but not AP2M1 phosphor-deficient mutant T156A in an *in vitro* kinase assay (Figure 2, B and C). Consistent with the higher kinase activity of LRRK2 G2019S, the AP2M1 T156 phosphorylation was two-fold greater with LRRK2 G2019S as compared to LRRK2 WT control (Figure 2, B and C). Moreover, the kinase dead LRRK2 did not induce AP2M1 phosphorylation in our *in vitro* kinase assay (Figure 2, B and C). These data therefore confirmed that LRRK2 phosphorylates AP2M1 at T156 *in vitro*.

**Figure 2.**
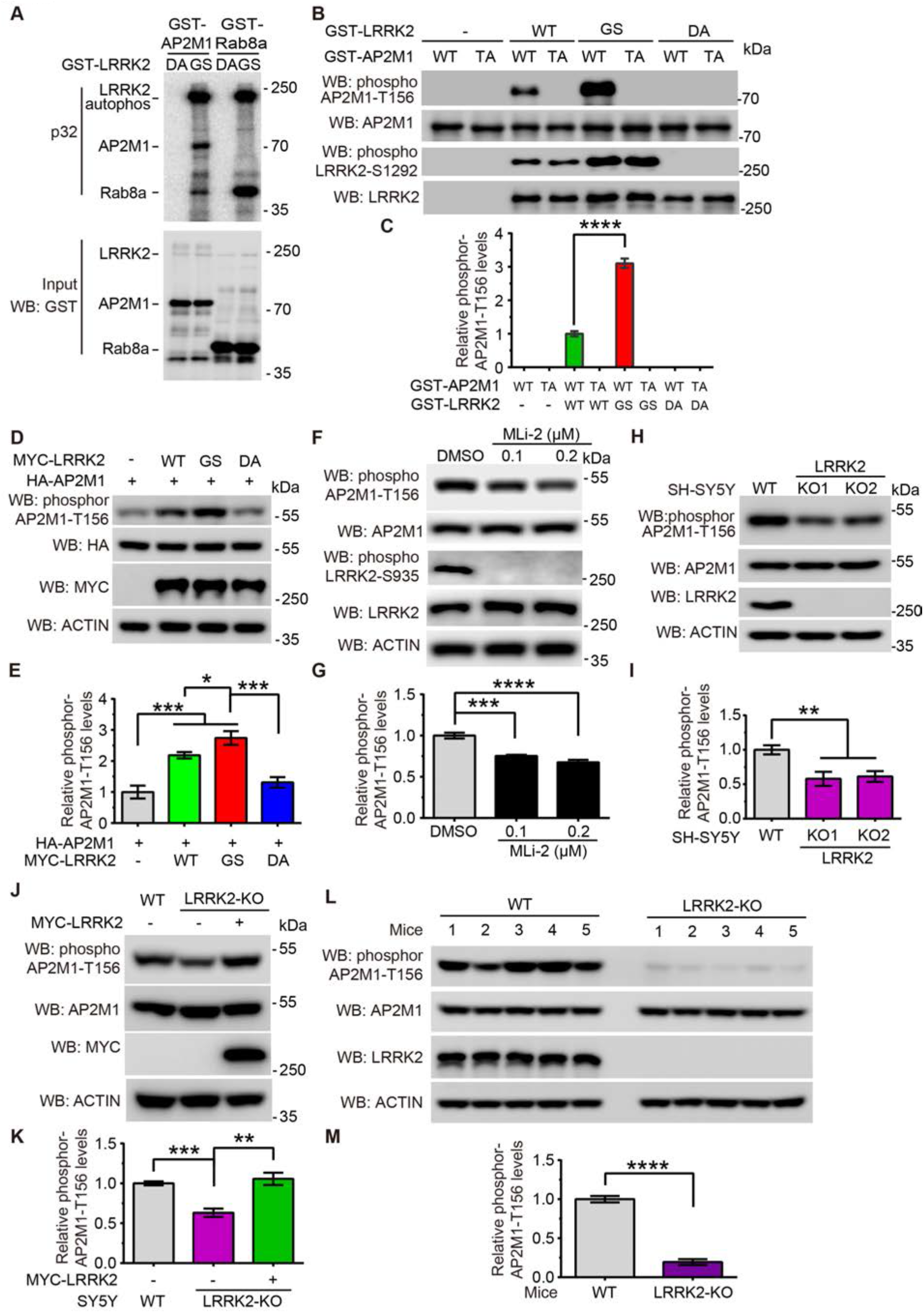
LRRK2 phosphorylates AP2M1 *in vitro* and *in vivo*. (**A**) LRRK2 phosphorylates AP2M1 *in vitro*. GST tagged G2019S (GS) and D1994A (DA) LRRK2 (970-2527aa) recombinant proteins were incubated with or without recombinant GST-AP2M1 and subjected to an *in vitro* kinase assay. Equivalent input is shown by GST immunoblotting for LRRK2 and AP2M1. Rab8a was as a positive control. (**B**) LRRK2 phosphorylates AP2M1 at T156 *in vitro*. GST tagged LRRK2 WT, GS and DA recombinant proteins were incubated with recombinant GST-AP2M1-WT, phosphor-deficient mutant T156A (TA) and subjected to an *in vitro* kinase assay followed by immunoblotting with AP2M1 phosphorylation specific antibody pT156. (**C**) Quantification of relative AP2M1 phosphorylation level at T156 by LRRK2, n=5. (**D**) Overexpression of LRRK2 WT and GS significantly increased AP2M1 cellular phosphorylation level but not LRRK2 kinase dead mutant DA. (**E**) Quantification of AP2M1 phosphorylation level. n=5. (**F**) Specific LRRK2 kinase inhibitor MLi-2 decreases endogenous AP2M1 phosphorylation levels. Lysates from MLi-2 treated SH-SY5Y cells at 0.1, 0.2 μM for 2 hours were subjected to immunoblotting with anti-pT156-AP2M1, anti-pS935-LRRK2. (**G**) Quantification of endogenous AP2M1 phosphorylation level upon MLi-2 inhibitor treatment. n=5. (**H**) LRRK2 phosphorylates endogenous AP2M1 *in vivo* in SH-SY5Y cells. AP2M1 has significantly lower phosphorylation levels in LRRK2 KO cells versus WT cells. (**I**) Quantification of AP2M1 phosphorylation level. n=5. (**J**) Overexpression of MYC-LRRK2 rescues the low level of AP2M1 phosphorylation in LRRK2 KO cells, suggesting a specificity of LRRK2 phosphorylation on AP2M1. Cell lysates form WT, LRRK2 KO, LRRK2 KO with overexpressed MYC-LRRK2 were subjected to immunoblotting with anti-pT156-AP2M1. (**K**) Quantification of AP2M1 phosphorylation level. n=5. (**L**) LRRK2 phosphorylates endogenous AP2M1 *in vivo* in mouse brains. AP2M1 has significantly lower phosphorylation levels in LRRK2 KO mouse brain lysates. (**M**) Quantification of AP2M1 phosphorylation level at T156 in WT, LRRK2 KO mouse brain lysates. n=5. Images were quantified by ImageJ software. Statistical significance was determined by Student’s *t* tests or One-way ANOVA followed by a Tukey’s post hoc test. Bar represents the mean ± SEM, \**P* < 0.05, \*\**P* < 0.01, \*\*\**P* < 0.001, \*\*\*\**P* < 0.0001.

To confirm our *in vitro* findings, we investigated the cellular phosphorylation of AP2M1 by overexpressing AP2M1 with hyperactive and kinase dead LRRK2, respectively, in SH-SY5Y cells. Consistent with our *in vitro* results, we found that LRRK2 hyperactive G2019S, but not LRRK2 kinase dead mutant D1994A, significantly increased AP2M1 cellular phosphorylation level (Figure 2, D and E). To further extend our findings to more physiological conditions, we examined the effect of LRRK2 on the phosphorylation of endogenous AP2M1. We first treated SH-SY5Y cells with MLi-2, a known specific LRRK2 kinase inhibitor, at different concentrations. As expected, MLi-2 largely inhibited LRRK2 kinase activity, as evidenced by abolished LRRK2 S935 phosphorylation. Consistently, endogenous AP2M1 T156 phosphorylation level decreased to about 70% upon MLi-2 treatment at 0.2 μM compared to the control (Figure 2, F and G), suggesting LRRK2 regulates AP2M1 phosphorylation at endogenous level. Next, we generated human neuroblastoma SH-SY5Y cell lines with depletion of LRRK2 using the CRISPR/Cas9 editing system. The resulting LRRK2 KO cells led to no detectable LRRK2 protein (Supplemental Figure 3, A and B). While endogenous AP2M1 T156 phosphorylation was detected in both WT and LRRK2 KO cells, the phosphorylation level was much lower in LRRK2 KO cells than in WT cells (Figure 2, H and I). To further confirm that the reduced AP2M1 T156 phosphorylation is specific due to LRRK2 knockout, we performed a rescue experiment by transfecting WT LRRK2 back into LRRK2 KO cells and examined the resulting AP2M1 phosphorylation. Indeed, we found that WT LRRK2 transfection completely recued AP2M1 phosphorylation to a level of about twofold than non-transfection LRRK2 KO cells (Figure 2, J and K), suggesting the specificity of LRRK2 on AP2M1 phosphorylation. Prompted by the above results, we next examined endogenous AP2M1 phosphorylation levels in both LRRK2 WT and KO mouse brains. We found that the levels of endogenous AP2M1 phosphorylation are significantly lower in LRRK2 KO brain lysate compared to WT brain lysate (Figure 2, L and M). Taken together, we concluded that LRRK2 phosphorylates AP2M1 at T156 both *in vitro* in cell cultures and *in vivo* in mouse brains.

### Tissue-specific role of LRRK2 on phosphorylation of AP2M1

It has been reported that AP2M1 is phosphorylated by a known kinase, AAK1 (AP2-associated protein kinase 1), at T156. We therefore investigated the regulatory relationship between LRRK2 and AAK1 on AP2M1 phosphorylation. To this end, we first performed *in vitro* kinase assay to compare the amount of ^32^P-ATP incorporated by AP2M1 upon equimolar amounts of LRRK2 and AAK1 (Figure 3, A and B). Our results showed that AAK1 is more potent than LRRK2 to phosphorylate AP2M1 *in vitro*. To ask whether LRRK2 and AAK1 have synergistic effects on AP2M1 phosphorylation, we examined AP2M1 phosphorylation levels treated with LRRK2 inhibitor MLi-2, or AAK1 inhibitor LP-935509, or a combination of MLi-2 and LP-935509. We found that while either MLi-2 or LP-935509 treatment reduces endogenous AP2M1 phosphorylation, treatment with both MLi-2 and LP-935509 has a much stronger effect on reducing AP2M1 phosphorylation (Figure 3, C and D), suggesting a synergistic effect between LRRK2 and AAK1 on AP2M1 phosphorylation. Based on above observed synergistic effect, we next asked whether the regulation of LRRK2 and AAK1 on AP2M1 depends on each other. To this end, we first overexpressed AAK1 in LRRK2 WT and KO cells, respectively, and determined the endogenous AP2M1 phosphorylation level. We found that while AAK1 can increase endogenous AP2M1 phosphorylation in both LRRK2 WT and KO cells, the increase of AP2M1 phosphorylation is more potent in LRRK2 KO cells than in LRRK2 WT cells (Figure 3, E and F), suggesting that AAK1 can phosphorylate AP2M1 independently of LRRK2. This was further confirmed by examining the effect of AAK1 on overexpressed AP2M1 phosphorylation in LRRK2 WT and KO cells, as we found that AAK1 phosphorylates overexpressed AP2M1 robustly in both LRRK2 WT and KO cells (Supplemental Figure 4, A and B). We noticed that LRRK2 KO cells exhibited a lower level of endogenous AP2M1 phosphorylation than WT cells with or without AAK1 overexpression, suggesting LRRK2 contributes a major part of endogenous AP2M1 phosphorylation (Figure 3, E and F). To investigate whether the effect of LRRK2 on AP2M1 phosphorylation relies on AAK1, we knocked down AAK1 expression in LRRK2 WT and KO cells. While the total phosphorylation level of AP2M1 is decreased upon AAK1 knockdown, AP2M1 phosphorylation in LRRK2-WT cells is more than that in LRRK2 KO cells, suggesting endogenous LRRK2 still phosphorylates AP2M1 despite AAK1 knockdown (Figure 3, G and H). This therefore suggests an independent regulation of AP2M1 phosphorylation by LRRK2. Notably, we failed to obtain AAK1 knockout cell lines even with multiple rounds of efforts using the CRISPR/Cas9 editing system, and shRNAs can only knock down half of AAK1, suggesting an essential role of AAK1 in SY5Y cells.

**Figure 3.**
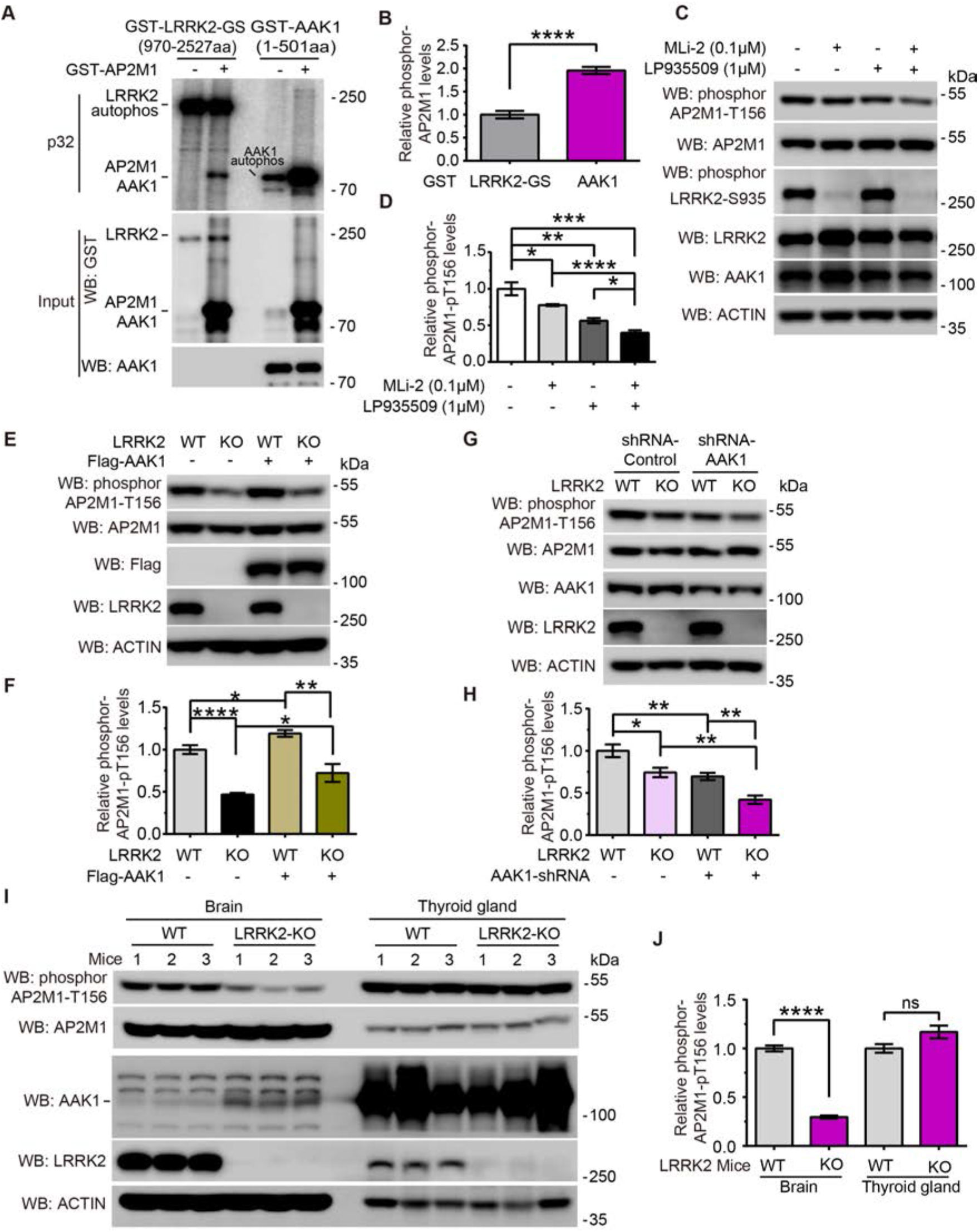
LRRK2 and AAK1 play an independent and tissue-specific role on phosphorylation of AP2M1. (**A**) The amount of ^32^P-ATP incorporated by AP2M1 upon equimolar amounts of GST-LRRK2-G2019S (GS) (970-2527aa) and GST-AAK1 (1-501aa) is measured by an *in vitro* kinase assay. GST-AAK1 (1-501aa) fragment and GST-AP2M1 have similar molecular weights, thus the signal in the line of GST-AAK1 with AP2M1 includes both AAK1 autophosphorylation and AP2M1 phosphorylation by AAK1. (**B**) Quantification of relative AP2M1 phosphorylation level at T156 by equimolar amounts of LRRK2 and AAK1, n=5. (**C**) LRRK2 inhibitor MLi-2 and AAK1 inhibitor LP-935509 synergistically decrease endogenous AP2M1 phosphorylation levels. Lysates from MLi-2 (0.1 μM) or/ and LP-935509 (1 μM) treated cells for 2 hours were subjected to immunoblotting with anti-pT156-AP2M1, anti-pS935-LRRK2. (**D**) Quantification of endogenous AP2M1 phosphorylation level upon MLi-2 and LP-935509 inhibitor treatment, n=5. (**E**) AAK1 phosphorylates AP2M1 independently of LRRK2. AAK1 was overexpressed in LRRK2 WT and KO cells. Lysates were subjected to immunoblotting with anti-pT156-AP2M1. (**F**) Quantification of endogenous AP2M1 phosphorylation level in LRRK2 WT and KO cells with overexpression of AAK1, n=5. (**G**) LRRK2 phosphorylates AP2M1 independently of AAK1. AAK1 was knocked down by shRNA to about 50% in LRRK2 WT and KO cells. Lysates were subjected to immunoblotting with anti-pT156-AP2M1. (**H**) Quantification of endogenous AP2M1 phosphorylation level in LRRK2 WT and KO cells with knockdown of AAK1, n=5. (**I**) LRRK2 plays a tissue-specific role on phosphorylation of AP2M1. AP2M1 has significantly lower phosphorylation levels in LRRK2 KO mouse brain lysates but not in LRRK2 KO mouse thyroid gland lysates. Lysates from brain and thyroid gland of WT and LRRK2 KO mice were subjected to immunoprecipitation with anti-pT156-AP2M1, LRRK2, AAK1. (**J**) Quantification of AP2M1 phosphorylation level at T156 in WT, LRRK2 KO mouse brain and thyroid gland lysates, n=5. Images were quantified by ImageJ software. Statistical significance was determined by Student’s *t* tests. Bar represents the mean ± SEM, \**P* < 0.05, \*\**P* < 0.01, \*\*\**P* < 0.001, \*\*\*\**P* < 0.0001.

We next asked the physiological relevance of AP2M1 phosphorylation by LRRK2 versus AAK1. LRRK2 is known for its high expression in the brain, where the PD pathologies are observed. Interestingly, we found that unlike LRRK2, the expression of AAK1 is relatively low in brain tissue (Figure 3, I and J). In contrast, AAK1 is highly expressed in the thyroid gland where LRRK2 expression is fairly low. The differential expression of LRRK2 and AAK1 observed in mouse brain versus thyroid gland prompted us to ask whether LRRK2 and AAK1 have tissue-specific roles in regulating AP2M1 phosphorylation. Indeed, knocking out LRRK2 in mouse brain dramatically decreased AP2M1 phosphorylation, suggesting LRRK2 plays a key role in regulating AP2M1 phosphorylation in brain. However, knocking out LRRK2 had no obvious effects on AP2M1 phosphorylation in thyroid gland where AAK1 is dominantly expressed, suggesting a minor role of LRRK2 on AP2M1 phosphorylation in thyroid gland (Figure 3, I and J). Interestingly, we noticed a mild increase of AAK1 protein level in LRRK2 knockout brains (Figure 3I), suggesting a feedback regulation between LRRK2 and AAK1 protein levels may exist. Taken together, our results suggest an AAK1-independent and tissue-specific role of LRRK2 on AP2M1 phosphorylation.

### LRRK2 phosphorylation of AP2M1 promotes AP2M1 membrane association

Having established a phosphorylation regulation of AP2M1 by LRRK2, we next asked the physiological function of this regulation. It is known that phosphorylation of AP2M1 enhances AP2 recruitment to the plasma membrane during coated pit assembly(16). We therefore asked whether phosphorylation of AP2M1 by LRRK2 regulates AP2M1 membrane association. To minimize the effect of endogenous LRRK2 on AP2M1, we overexpressed MYC-LRRK2 and HA-AP2M1 in LRRK2 KO SY5Y cells, and subsequently fractionated these cells into cytosol and membrane fractions and determined the abundance of soluble or membrane-associated AP2M1 by Western blot (WB). AP2M1 phosphor-deficient mutant AP2M1-T156A was introduced as a control. Our results showed that LRRK2-GS, the kinase active mutant, potentiated AP2M1-WT membrane localization but not AP2M1-T156A mutant (Figure 4, A and B). To further confirm this finding, we employed an established liquid nitrogen coverslip freeze-thaw method(17, 18). Using this method, we were able to deplete cytosolic proteins and fix membrane fraction on the coverslip, allowing us to further examine the localization of membrane-associated AP2M1. LRRK2 KO SY5Y cells were transfected with DsRed-LRRK2 WT, GS or kinase dead mutant DA and transduced with lentiviruses carrying eGFP-AP2M1-WT or T156A. 48 hours after transfection, cells were permeabilized by liquid nitrogen freeze–thaw to deplete cytosol and then fixed and visualized by confocal microscopy. Membrane associated AP2M1-WT, but not AP2M1-T156A, was significantly increased with LRRK2 WT or GS co-expression (Figure 4, C and E). The effect of LRRK2 on AP2M1 membrane localization was further confirmed by TIRF (Total Internal Reflection Fluorescence) microscopy (Figure 4, D and F). Consistently, LRRK2 WT and GS significantly increased AP2M1-WT but not AP2M1 - T156A membrane association (Figure 4, D and F, Supplemental Figure 5). Together, these results suggest that LRRK2 phosphorylation of AP2M1 promotes AP2M1 membrane association.

**Figure 4.**
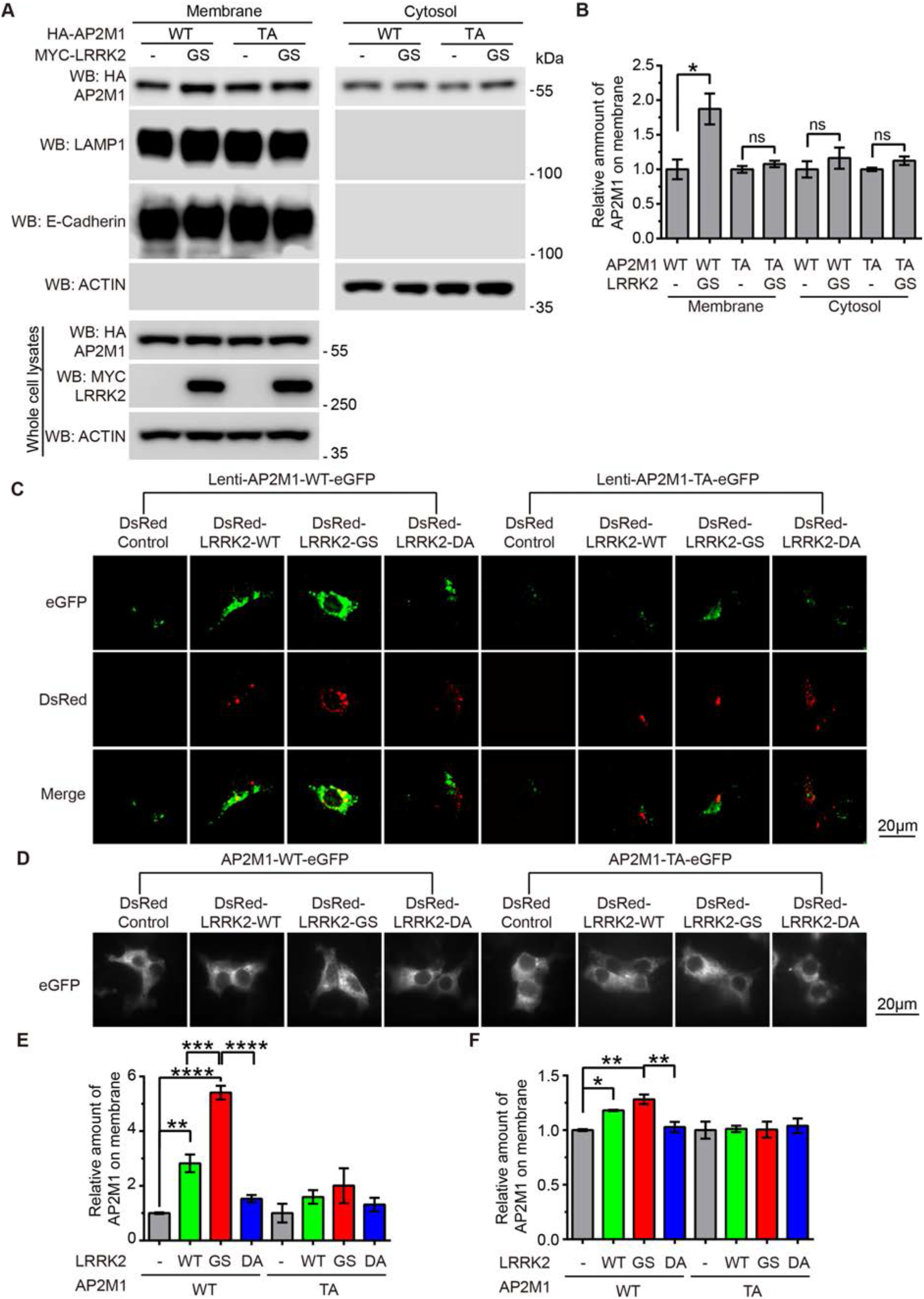
LRRK2 phosphorylation of AP2M1 increases AP2M1 membrane association. (**A**) LRRK2 GS increases AP2M1-WT membrane association but not AP2M1-T156A by cellular fractionation. LRRK2 KO SH-SY5Y cells were transfected with HA-AP2M1-WT or T156A (TA) with or without MYC-LRRK2-GS. After 48 hours, cells were harvested and fractionated into cytosol and membrane fractions. (**B**) Quantification of AP2M1 intensity on membrane and cytosol fractions. Images were quantified by ImageJ software. Statistical significance was determined by Student’s *t* tests. Bar represents the mean ± SEM, n=5, \**P* < 0.05. (**C**) LRRK2 increases AP2M1 membrane association by liquid nitrogen coverslip freeze-thaw methods. LRRK2 KO SH-SY5Y cells were transfected with DsRed-LRRK2 WT, GS, or DA and transduced with lentiviruses carrying eGFP-AP2M1 WT or TA. After 48 hours, cells were permeabilized by liquid nitrogen freeze–thaw to deplete cytosol and then fixed and visualized by confocal microscopy. Scale = 20 μm. (**D**) LRRK2 increases AP2M1 membrane association by TIRF microscopy. LRRK2 KO SH-SY5Y cells were cotransfected with DsRed-LRRK2 WT, GS, or DA and eGFP-AP2M1 WT or TA at a plasmid ratio of 5:1. After 48 hours, cells were fixed and imaged by TIRF microscopy. Scale = 20 μm. (**E**) Quantification of AP2M1 intensity on membrane by liquid nitrogen coverslip freeze–thaw methods. (**F**) Quantification of AP2M1 intensity on membrane by TIRF microscopy. Images were quantified by ImageJ software. Statistical significance was determined by one-way ANOVA followed by a Tukey’s post hoc test. Data represent the mean ± SEM from three independent experiments with n ≥ 10 cells quantified in each experiment, \**P*<0.05, \*\**P*<0.01, \*\*\**P*<0.001, \*\*\*\**P*<0.0001.

### LRRK2 phosphorylation of AP2M1 mediates LRRK2-induced endocytosis defects in neurons

Next, we asked what is the functional consequence of increased AP2M1 membrane association by LRRK2 phosphorylation. AP2M1 phosphorylation specifically enhances its association with cargos such as transferrin (Tfn) receptor, which in turn, mediates endocytosis (19, 20). Therefore, we monitored endocytic trafficking using Tfn-594 dye internalization. To study in a more physiological condition, we examined the endocytosis in wildtype, LRRK2 KO and G2019S knockin (GSKI) neurons by monitoring Tfn internalization. Surface Tfn was removed by acid stripping. The internalized Tfn in neuronal cell body was monitored (Figure 5). Overexpression of AP2M1-WT or T156A in wildtype neurons induced decreased Tfn internalization, indicating a defect in endocytosis. Both LRRK2 GSKI and KO neurons also induced defects in Tfn internalization (Figure 5). Interestingly, Tfn internalization defects in LRRK2 GSKI neurons were enhanced by overexpression of AP2M1-WT but not AP2M1-TA mutant (Figure 5, A and C). However, no significant difference was observed when overexpression of AP2M1-WT or T156A in LRRK2 KO neurons (Figure 5, A and C). This suggests that increased AP2M1 phosphorylation by LRRK2 induces more severe defects in endocytosis. To further examine if LRRK2 phosphorylation impairs AP2M mediated endocytosis, we inhibited endogenous LRRK2 kinase activity in primary neurons by MLi-2 treatment and examined the subsequent endocytosis of neurons overexpressing AP2M1-WT or T156A (Figure 5, B and D). Indeed, inhibition of LRRK2 kinase rescued AP2M1-WT mediated endocytic defects (Figure 5, B and D). Together, these data suggest LRRK2 phosphorylation of AP2M1 mediates LRRK2-induced endocytosis defects in neurons.

**Figure 5.**
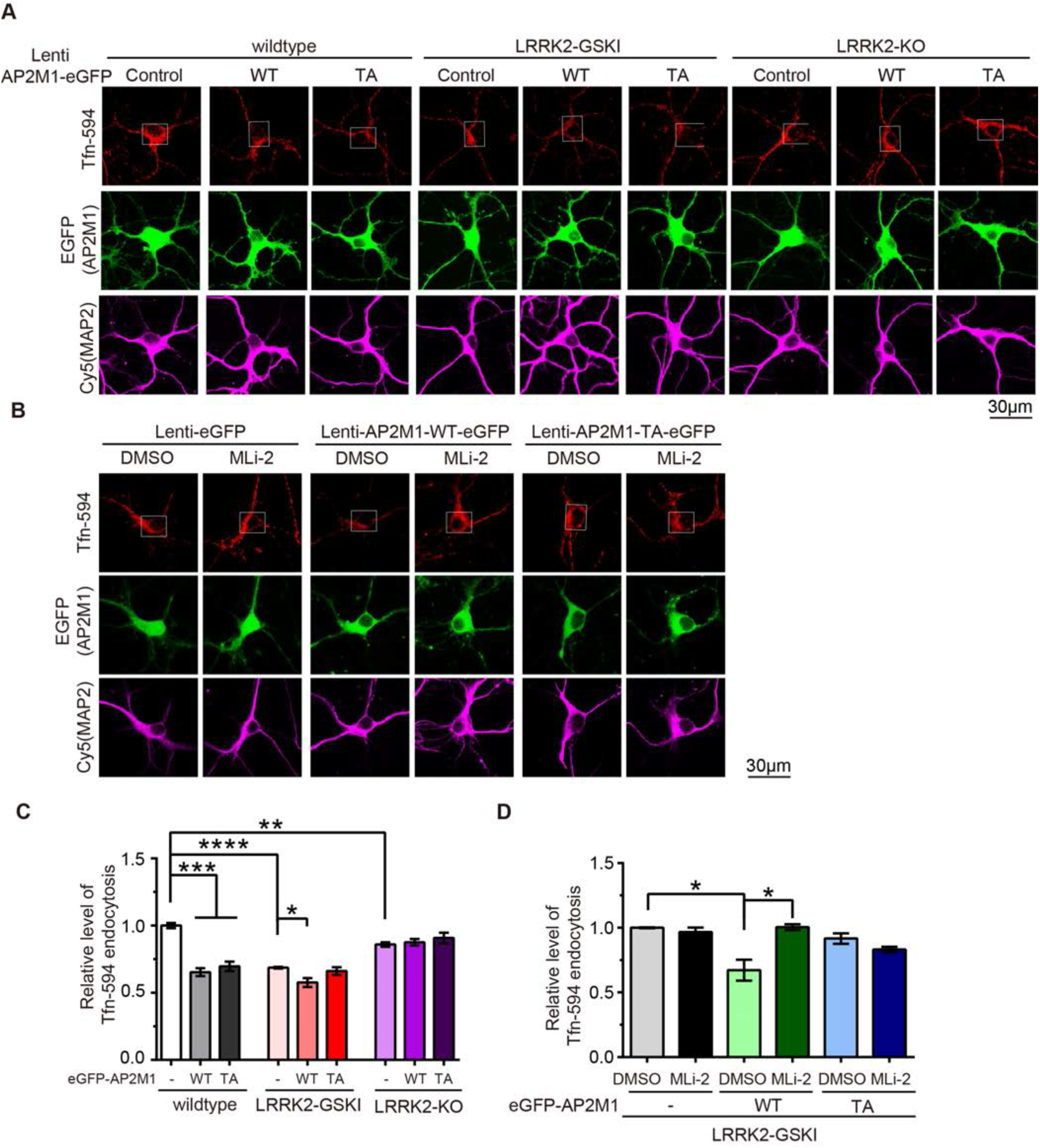
LRRK2 phosphorylation of AP2M1 mediates LRRK2 induced endocytosis defects in neurons. (**A**) Lentiviruses carrying eGFP-AP2M1 WT or T156A (TA) were transduced into LRRK2 WT, KO or GSKI hippocampal neurons and Tfn internalization was monitored. (**B**) Lentiviruses carrying eGFP-AP2M1 WT or T156A (TA) were transduced into LRRK2 GSKI hippocampal neurons. 0.1 μM MLi-2 was pre-incubated for 2 hours in LRRK2 GSKI hippocampal neurons carrying AP2M1-WT or TA lentiviruses. Tfn internalization was monitored. Scale = 30 μm. (**C**) Quantification of **A** Tfn internalization in at the cell bodies (indicated in white boxes) of LRRK2 WT, KO or GSKI hippocampal neurons transduced with eGFP-AP2M1 WT or TA. d Quantification of **B** Tfn internalization at the cell bodies. Statistical significance was determined by Student’s *t* tests or one-way ANOVA followed by a Tukey’s post hoc test. Data are the means ± SEM. Three independent experiments with n ≥ 15 cells quantified in each experiment. \**P* < 0.05, \*\**P* < 0.01, \*\*\**P* < 0.001, \*\*\*\**P* < 0.0001.

### Excessive LRRK2 kinase activity inhibits AP2M1 uncoating

How do both LRRK2 KO and GSKI induce endocytic defects? One explanation could be that LRRK2 regulates AP2M1 phosphorylation and dephosphorylation cycles. Phosphorylation of AP2M1 enhances AP2 recruitment to the plasma membrane during initial coated pit assembly(16). After coated pits are assembled, coated pits will be scissored and endocytosed into cytosol. Following this process, AP2M1 needs to be dephosphorylated and uncoated from the pits, and then recycled back to the plasma membrane for a new cycle of coated pit endocytosis (13) (Figure 9C). LRRK2 KO could impair basal level of AP2M1 phosphorylation for initial coated pit assembly while excessive LRRK2 kinase activity could inhibit AP2M1 dephosphorylation, which in turn may inhibit AP2M1 uncoating from the pits (Figure 9C). To test this possibility, we examined the effects of LRRK2 GS and DA on AP2M1 uncoating using an *in vitro* uncoating assay (21). Purified CCVs from mouse brains (Figure 6, A-C) were incubated with purified GST-LRRK2 GS or DA in the presence of an ATP-regenerating system, GST protein used as a control. AP2M1 staying on CCV pellets was analyzed by WB. The amount of AP2M1 on CCV indicates how much of AP2M1 was not successfully uncoated from CCVs and released into supernatant during uncoating. LRRK2 GS retained more AP2M1 on CCV pellets (Figure 6, D and E), indicating an inhibition of AP2M1 uncoating. These data are consistent with our previous study by transmission electron microscopy (TEM) showing that specific expression of LRRK2 GS in dopaminergic neurons in mouse brain induces a robust reduction in the number of synaptic vesicles and an accumulation of CCVs at synapses *in vivo* (8).

**Figure 6.**
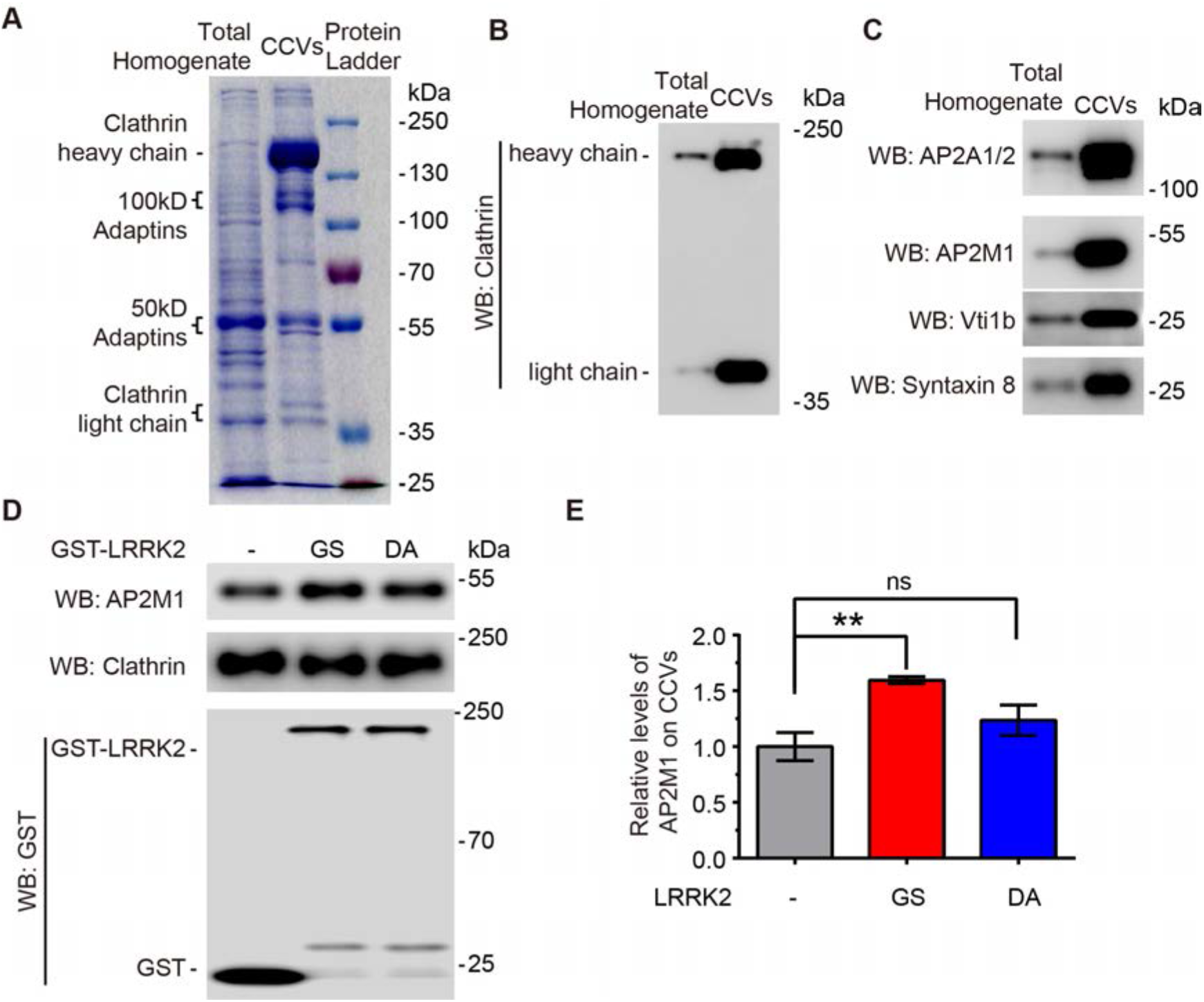
Excessive LRRK2 kinase activity inhibits AP2M1 uncoating. (**A**) Protein profile of the total homogenate and CCVs purified from mouse brains after SDS-PAGE and Coomassie blue staining. (**B and C**) Characterization of purified CCVs by immunoblotting with anti-Clathrin heavy chain and light chain **B** and with antibodies of CCV enriched proteins as indicated **C**. (**D**) Purified CCVs from mouse brains were incubated with purified GST-LRRK2 GS or DA in the presence of an ATP-regenerating system. AP2M1 staying on CCV pellets, which indicates uncoated AP2M1 from CCVs, was analysis by Western blot. (**E**) Quantification of AP2M1 staying on CCVs. Images was quantified by ImageJ software. Statistical significance was determined by one-way ANOVA followed by a Tukey’s post hoc test. Data are the means ± SEM, n = 5, \*\**P* < 0.01.

### LRRK2 phosphorylation of AP2M1 mediates LRRK2-induced neuronal toxicity

To ascertain whether LRRK2 phosphorylation of AP2M1 mediates LRRK2-induced neuronal toxicity, the effects of AP2M1 WT and T156A on LRRK2 WT, GS, DA-induced toxicity were examined. AP2M1 and LRRK2 were transfected into primary cortical neuron cultures and the toxicity was characterized by neurite loss and cell death(15, 22–25). LRRK2 WT and GS both caused neuronal toxicity while LRRK2 GS induced toxicity to a more severe content than LRRK2 WT, which is consistent as previous reports(15, 22–25). Overexpression of AP2M1 WT or T156A alone also induced neuronal toxicity. Interestingly, overexpression of AP2M1 WT significantly enhanced LRRK2 WT or GS-induced neuronal toxicity while overexpression of AP2M1 T156A had little effect (Figure 7, A and B). This was further confirmed by TUNEL staining (Terminal deoxynucleotidyl transferase dUTP nick end labeling) (Supplemental Figure 6). These data suggest that LRRK2 phosphorylation of AP2M1 mediates LRRK2-induced neuronal toxicity.

**Figure 7.**
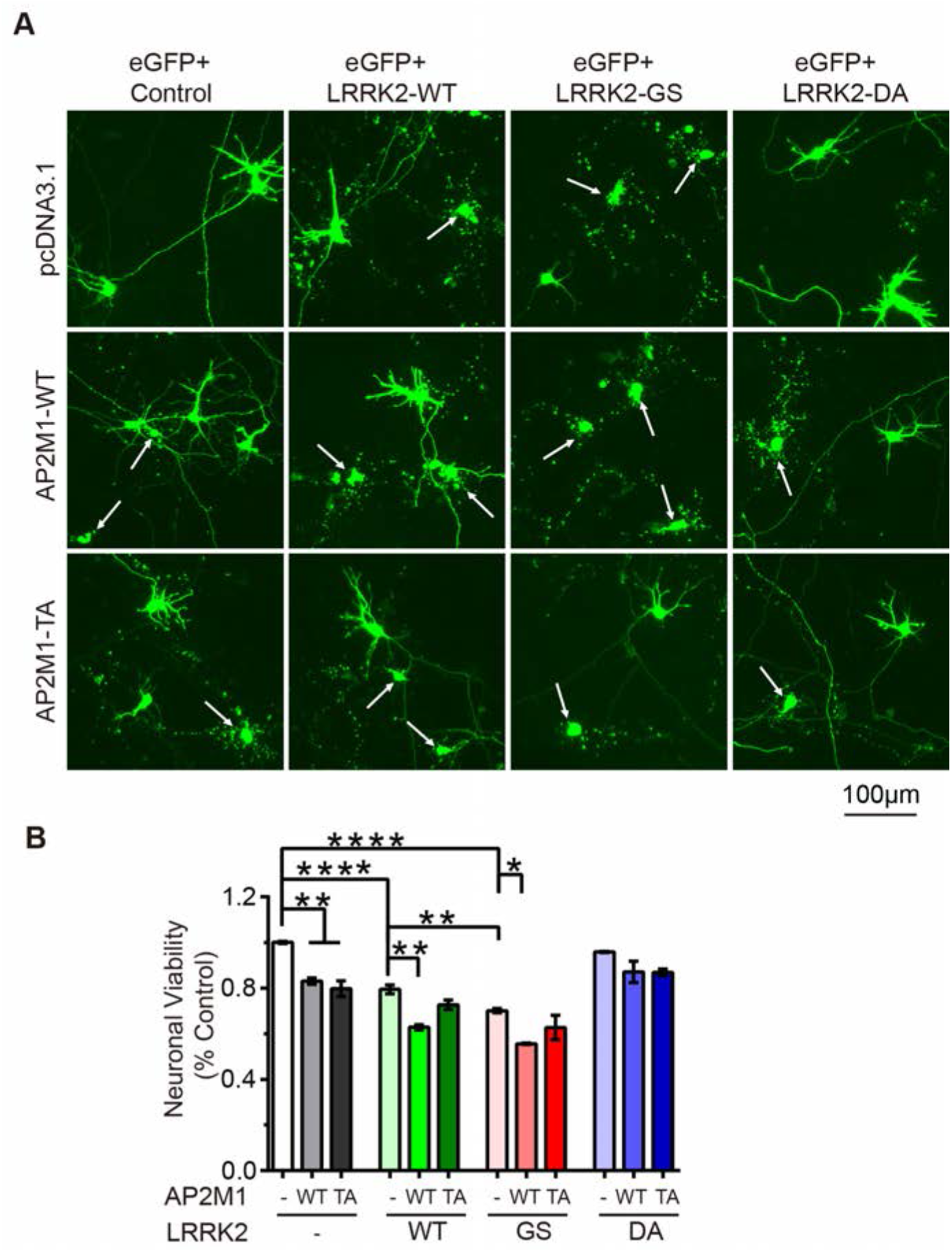
LRRK2 phosphorylation of AP2M1 mediates LRRK2-induced neuronal toxicity. (**A**) AP2M1 WT enhances LRRK2 induced neuronal toxicity. Representative fluorescent images (green) showing that mouse primary cortical neurons were co-transfected with MYC-LRRK2, HA-AP2M1 with eGFP at a plasmid ratio of 10:10:1 at DIV 5. Neuronal viability was analyzed at 48-hour post-transfection with non-viable neurons exhibiting no obvious neurite process (arrow). Scale = 100 μm. (**B**) Quantification of neuronal viability. Bars indicate the viability (n > 100) for each transfection condition expressed as a percent of control neurons (eGFP with pcDNA3.1 empty vector). Statistical significance was determined by one-way ANOVA followed by a Tukey’s post hoc test. Data represent the mean ± SEM from three independent experiments with n > 100 cells quantified in each experiment. \**P* < 0.05, \*\**P* < 0.01, \*\*\*\**P* < 0.0001.

### LRRK2 regulated AP2M1 phosphorylation modulates LRRK2-induced dopaminergic neurodegeneration *in vivo*

To determine whether AP2M1 phosphorylation by LRRK2 modulates LRRK2 dopaminergic neurodegeneration *in vivo*, we used LRRK2 *Drosophila* model in which flies express human LRRK2-WT or GS. We also generated transgenic flies carrying UAS-AP2M1-WT and T156A. The transgenic flies were crossed with *dopa decarboxylase (Ddc)-Gal4>UAS-GFP* driver flies to achieve specific expression of transgene in dopaminergic and serotonin neurons. Dopaminergic neuron number as revealed by GFP fluorescence was monitored in four major dopaminergic neuronal clusters in *Drosophila* adult brain (PPM1/2, PPM3, PPL1) (Figure 8, A and B)(26). Consistent as previous LRRK2 *Drosophila* models(27), our LRRK2 WT, GS flies exhibit substantial dopaminergic neurodegeneration in the major dopaminergic clusters (Figure 8, C-E) and total dopaminergic neurons (Figure 8F). Overexpression of AP2M1-WT or T156A alone also induced dopaminergic neurodegeneration in the major dopaminergic clusters (Figure 8, C-E) and total dopaminergic neurons (Figure 8F). Notably, AP2M1-WT enhanced both LRRK2-WT and GS induced dopaminergic neurodegeneration as indicated by the decreased total dopaminergic neuron numbers while AP2M1-T156A had little effect (Figure 8F), which is consistent with our neuronal culture results.

**Figure 8.**
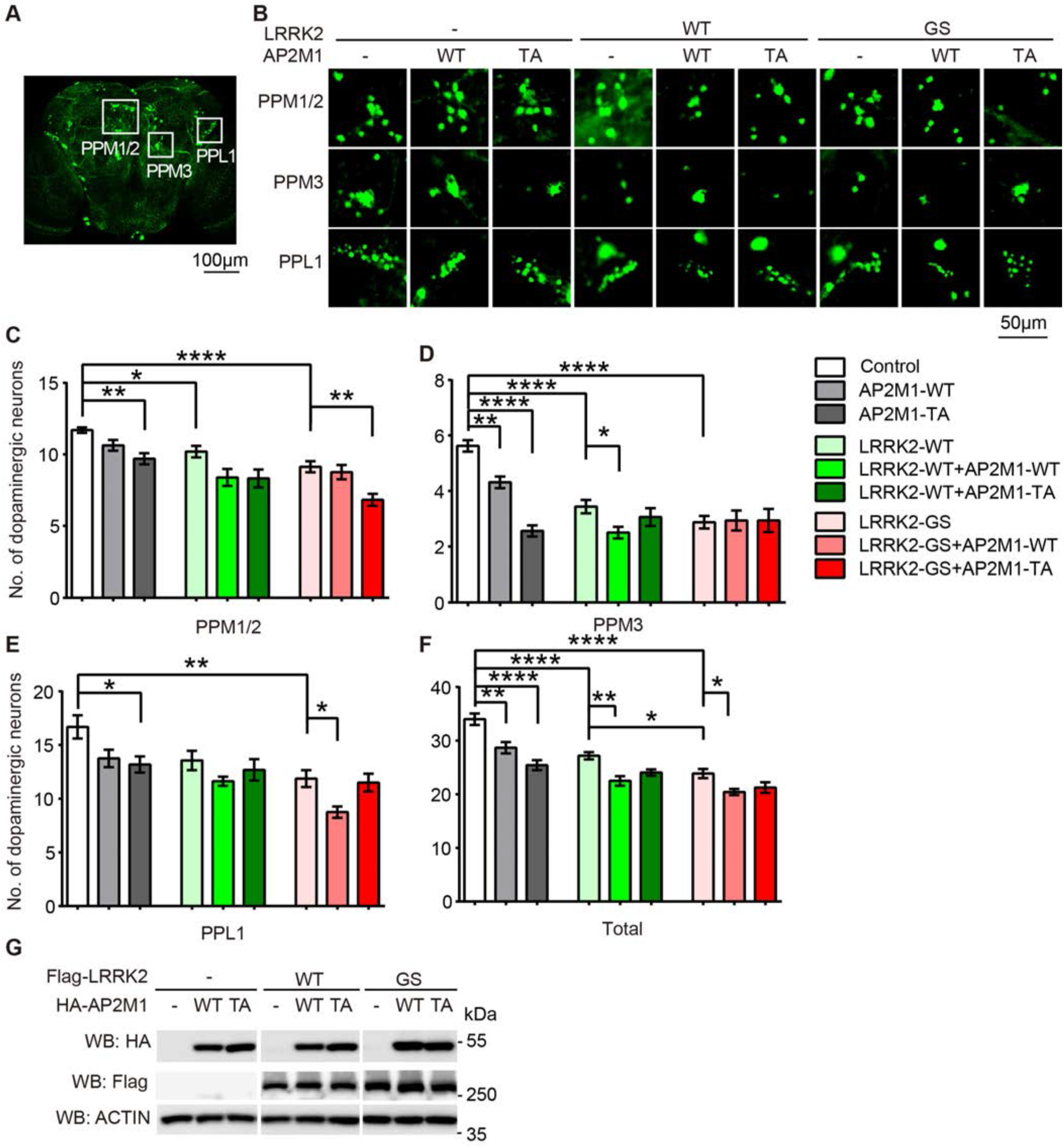
LRRK2 phosphorylation of AP2M1 modulates LRRK2-induced dopaminergic neurodegeneration *in vivo*. (**A**) Diagram of dopaminergic neuronal clusters (PPM1/2, PPM3, PPL1) in the posterior areas of the adult fly brain. Scale = 100 μm. (**B**) Representative confocal images (GFP) of dopamine neurons in each dopaminergic cluster from 7-weeks old flies of the indicated genotypes. Scale = 50 μm. (**C-E**) Quantification of dopamine neurons per dopaminergic cluster in 7-weeks old flies of the indicated genotypes. (**F**) Total numbers of dopamine neurons in four major dopaminergic clusters of the flies at 7-weeks old. Statistical significance was determined by one-way ANOVA followed by a Tukey’s post hoc test. Data are the means ± SEM, n = 8 flies per genotype, \**P* < 0.05, \*\**P* < 0.01, \*\*\**P* < 0.001, \*\*\*\**P* < 0.0001. (**G**) Expression levels of overexpressed LRRK2 and AP2M1 in flies driven by GMR-Gal4. Lysates prepared form whole heads of one-week old flies of the indicated genotypes were subjected to immunoblotting with anti-HA-HRP, anti-Flag-HRP and anti-fly-actin.

### AP2M1 phosphorylation levels are elevated in the fibroblasts of both LRRK2-assocaited PD and sporadic PD

To test the potential pathophysiologic relevance of AP2M1 phosphorylation by LRRK2 in PD pathogenesis, we monitored AP2M1 phosphorylation levels and LRRK2 levels in human fibroblasts from PD patients carrying LRRK2 GS mutation and sporadic PD and age-matched health controls (Figure 9, A and B). The AP2M1 T156 phosphorylation levels are increased in LRRK2 GS mutation carrying fibroblasts and sporadic PD fibroblasts (Figure 9, A and B). This data suggests AP2M1 phosphorylation at T156 may play a role in the pathology of LRRK2-associated PD and sporadic PD.

**Figure 9.**
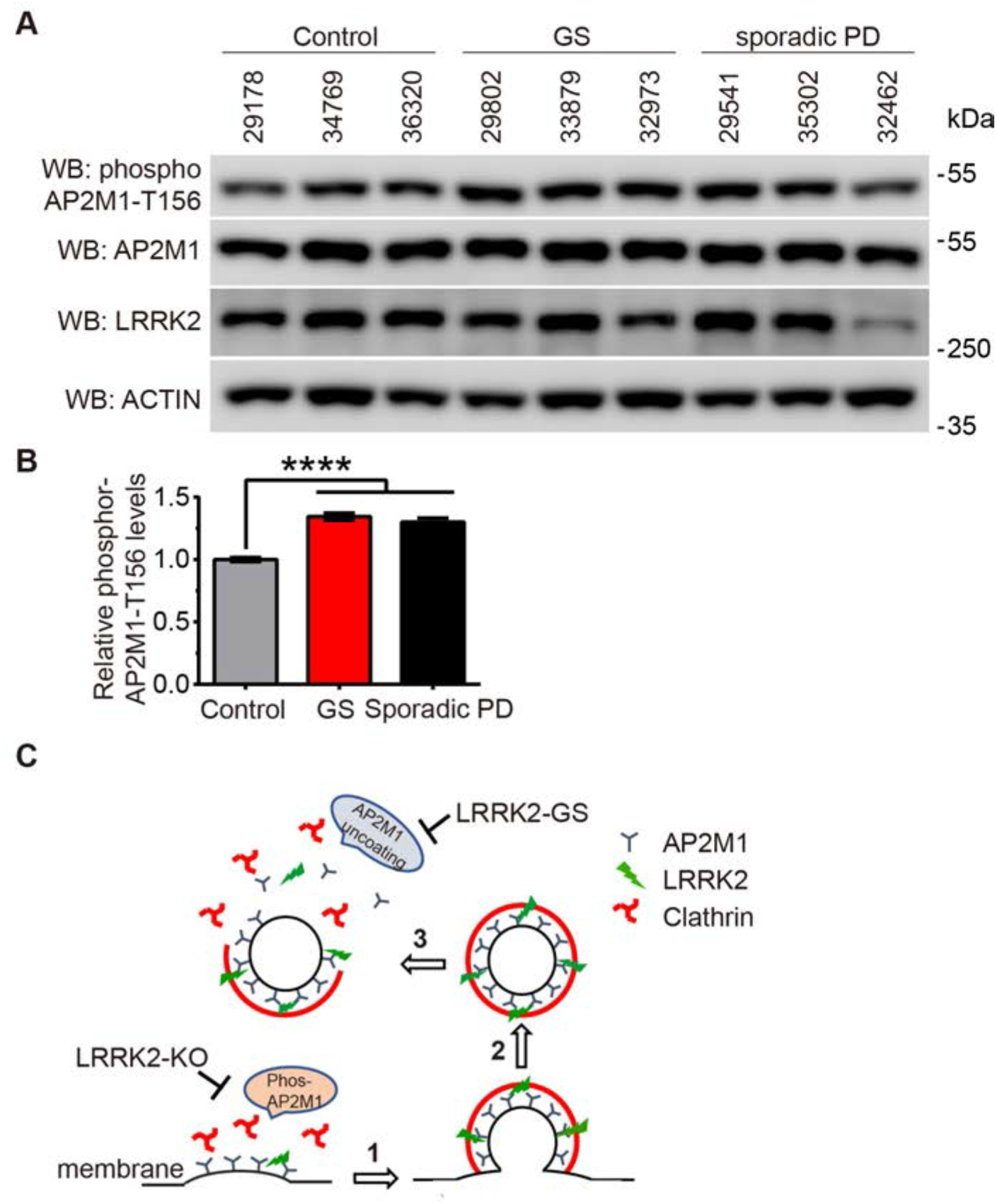
AP2M1 phosphorylation levels are elevated in the fibroblasts of LRRK2-assocaited PD and sporadic PD patients. (**A**) Levels of pT156-AP2M1, total AP2M1 and LRRK2 in PD patient fibroblasts. Lysates prepared form patient fibroblasts of the indicated LRRK2 GS and sporadic PD were subjected to immunoblotting with anti-pT156-AP2M1, anti-AP2M1, anti-LRRK2, and anti-actin. (**B**) Quantification of pT156-AP2M1 levels in PD patient fibroblasts. Images were quantified with ImageJ software. Statistical significance was determined by one-way ANOVA followed by a Tukey’s post hoc test. Data are the means ± SEM, n = 5, \**P* < 0.05. (**C**) Model of LRRK2 regulation on AP2M1 phosphorylation cycles to mediate endocytosis. Endogenous LRRK2 phosphorylates AP2M1 at T156 at plasma membrane, increases AP2M1 membrane association and in turn promotes the initiation of CCV formation (step **1**). After CCV scission, AP2M1 dephosphorylation promotes its uncoating from CCVs, a critical process required for the new cycle of CCV formation (step **3**). However, overexpression of LRRK2 or excessive LRRK2 kinase activity such as LRRK2 GS mutant inhibits AP2M1 dephosphorylation after CCV scission and results in inhibition of AP2M1 uncoating for the new cycle of CCV formation.

## Discussion

The major finding of this study is that LRRK2 interacts with and directly regulates the phosphorylation cycle of AP2M1, one of the major coat constituents in CME and recently implicated in PD risk(2), leading to the dysregulation of CME and dopaminergic neurodegeneration. We demonstrated that either knockout or excessive LRRK2 kinase activity induces endocytic defects, which is caused by LRRK2-induced abnormal AP2M1 phosphorylation cycle. These findings are consistent with a model in which knockout of LRRK2 decreases the basal level of AP2M1 phosphorylation (Figure 9C, step 1), which is required for the initial CCV formation, while excessive LRRK2 kinase activity inhibits AP2M1 dephosphorylation after CCV scission and results in inhibition of AP2M1 uncoating for the new cycle of CCV formation (Figure 9C, step 3). Further, AP2M1-WT but not T156A mutant enhanced LRRK2 WT and GS induced neuronal toxicity *in vitro*, and dopaminergic neurodegeneration *in vivo*. This suggests that LRRK2 phosphorylation of AP2M1 mediates LRRK2-induced neuronal toxicity both *in vitro* and *in vivo*. Importantly, AP2M1 phosphorylation levels are elevated in the fibroblasts of both LRRK2-assocaited PD and sporadic PD patients. This indicates that AP2M1 phosphorylation by LRRK2 plays an important role in modulation of mutant LRRK2 pathogenesis, and its’ involvement in sporadic PD pathology.

A recent systematic analysis of pathway-specific genetic risk factors revealed that the endocytic membrane trafficking pathway plays a major role in the risk of PD(2). Interestingly, *AP2M1* is one of the risk genes among the most contributors of endosomal membrane-trafficking pathway to the risk of PD(2). However, the biological process and the molecular mechanism underlying how AP2M1 involves in neurodegeneration and PD pathogenesis is unknown. Our study provides an attractive mechanistic link that AP2M1-mediating endocytosis is regulated through phosphorylation at T156 by LRRK2, the most common mutated protein of both sporadic and familial PD, leading to dopaminergic neurodegeneration. Interestingly, we and others previously reported that ArfGAP1 acts as a GTPase activating protein (GAP) and also a kinase substrate for LRRK2 to regulate neuronal toxicity *in vitro* and *in vivo*(15, 28). ArfGAP1 was reported to promote AP2-dependent endocytosis (29). It would be interesting to test whether LRRK2, ArfGAP1 and AP2 form a protein complex to regulate vesicular endocytosis and mediate neurodegeneration.

Mutations in LRRK2 have been demonstrated to cause defects in endocytosis(4, 5). Specific expression of the most prevalent LRRK2 mutation GS in dopaminergic neurons in mouse brain induces a robust reduction in the number of synaptic vesicle and an accumulation of CCVs at synapses, indicating a CME trafficking defect *in vivo*(8). But how LRRK2 mediates endocytosis is largely unknown. LRRK2 has been reported to interact with and phosphorylate several endocytic proteins including Endophilin A1 (EndoA)(30), synaptojanin (SYNJ1) (31, 32), Auxilin (DNAJC6) (33), and Rabs (17, 34–38). These proteins, including AP2M1, perform distinct functions at different steps in endocytic pathway, raising the possibility that LRRK2 acts as a regulatory kinase for multiple proteins at endocytic trafficking and LRRK2-induced neuronal toxicity could a result of a combination of several aspects of endocytic trafficking defects. Interestingly, SYNJ1, Auxilin, AP2M1 are also risk factors for PD and are linked to CME (2, 39, 40).

LRRK2 phosphorylates AP2M1 at T156, the site that has been reported to be phosphorylated by a known kinase, adaptor-associated kinase 1 (AAK1)(41–43). Interestingly, our study suggests that LRRK2 and AAK1 have independent roles on AP2M1 T156 phosphorylation. Importantly, our study also revealed a tissue-specific role of LRRK2 on AP2M1 phosphorylation, suggesting potential pathophysiologic relevance of AP2M1 phosphorylation by LRRK2. Notably, a single nucleotide polymorphism in an intron of the AAK1 gene has been associated with the age of onset of PD(44, 45), further suggesting impaired AP2M1 phosphorylation at T156 may be associated with PD, and maintaining proper phosphorylation cycle of AP2M1 may play an important role in PD pathogenesis.

Collectively, we show that LRRK2 acts as a kinase for AP2M1 and regulates the phosphorylation cycle of AP2M1, a major player in CME and a newly implicated factor in PD risk, leading to defects in CME and in turn the neuronal toxicity and dopaminergic neurodegeneration. Our study further supports the importance of endocytic pathway in PD pathology.

## Materials and Methods

### Animals

LRRK2 GSKI mice were purchased from Taconic (13940)(46) and LRRK2 KO mice were purchased from JAX Laboratory (16121)(47). Mice were housed and treated in accordance with the National Institutes of Health (NIH) ‘Guide for the Care and Use of Laboratory Animals’ and Institutional Animal Care and Use Committees of Kansas State University. Animals were housed in a 12-hour dark and light cycle with free access to water and food.

### Plasmids

Entry clones carrying AP2A2, AP2B1, AP2M1, AP2S1, AAK1 full-length cDNA were obtained from DNASU (clones HsCD00042465, HsCD00042299, HsCD00041654, HsCD00399704, HsCD00718566). Full-length human AP2A2, AP2B1, AP2M1, and AP2S1 were cloned by Gateway technology into the mammalian expression vector pcDNA3.1-3HA or 3flag-DEST, which was generating by replacing a V5 tag with a 3HA tag or a 3flag tag from pcDNA3.1-nV5-DEST vector (Invitrogen), and *E. coli* expression vector pDEST15 with GST tag Gateway Vector (Invitrogen). MYC-LRRK2-WT was a gift from Dr. Ted Dawson (Addgene plasmid # 17609, http://n2t.net/addgene:17609; RRID: Addgene_17609)(48). pENTR221 LRRK2 WT was a gift from Michael J Fox Foundation MJFF (Addgene plasmid # 39529). Full-length human LRRK2 and AP2M1, were cloned by Gateway technology into the mammalian expression vector pcDNA3.1-eGFP-DEST or pcDNA3.1-DsRed-DEST, which was generating by replacing a V5 tag with an eGFP or DsRed (Clontech) tag from pcDNA3.1-nV5-DEST vector (Invitrogen). To generate AP2M1 or LRRK2 single or multiple mutants, site-directed mutagenesis were carried out using the In-Fusion HD Cloning Plus (Clontech). AP2M1 domain fragments (D1, D2, D3, D4) were cloned into pcDNA3.1 vector with a N-terminal HA tag. Truncated mutants for human LRRK2 were cloned into the mammalian expression vector pcDNA3.1 with three flag tags at N-terminus as described previously(15). Full-length AP2M1 WT and T156A were cloned into *Drosophila* Gateway expression vector pUAST vector with an N-terminal HA tag (Drosophila Genomics Resource Center, Bloomington, IN). Full-length AP2M1 WT and T156A were cloned into pFUGW lentiviral vector with an C-terminal eGFP tag. All cDNAs or mutation sites were confirmed by DNA sequencing analysis.

### Antibodies and reagents

Rabbit monoclonal anti-AP2M1 (Ab759976), rabbit monoclonal anti-p-T156 AP2M1 (ab109397), rabbit monoclonal anti-p-S1292-LRRK2 (ab203181), rabbit polyclonal anti-GFP, rabbit polyclonal anti-GST-HRP (ab3416) antibodies were obtained from Abcam. Mouse monoclonal anti-LRRK2 (clone N138/6) was from UC Davis/NIH NeuroMab facility. Rabbit monoclonal anti-Clathrin Heavy Chain (CHC, 4796) and mouse monoclonal anti-E-Cadherin (4A2, 14472) were obtained from Cell Signalling Technology. Mouse monoclonal anti-AAK1 was obtained from Santa Cruz Biotechnology (sc-134242). Rabbit polyclonal anti-Syntaxin 8 (110083) and anti-Vti1b (164002) were obtained from Synaptic Systems. Mouse monoclonal anti-LAMP1 was obtained from Developmental Studies Hybridoma Bank (clone H4A3, DSHB). Mouse monoclonal anti-V5 antibody (R96025) was purchased from Thermo fisher. Mouse monoclonal anti-MYC, anti-V5-HRP, anti-flag, anti-flag-HRP, anti-HA, anti-HA-HRP, and anti-actin antibodies were obtained from Sigma-Aldrich. HRP-linked anti-rabbit or anti-mouse IgG antibodies were obtained from Jackson ImmunoResearch Labs. AlexaFluor-488 anti-mouse IgG, AlexaFluor-488 anti-rabbit IgG, Cyanine5 (CY5) anti-mouse IgG antibodies, and AlexaFluor 594-conjugated Tfn were obtained from Molecular Probes (Thermofisher).

### PD patient fibroblasts

The fibroblasts from health controls (ND34769, ND29178, ND36320), from patients carrying LRRK2 G2019S mutation (ND33879, ND29802, ND32973), from sporadic PD (ND35302, ND29541, ND32462) were obtained from the NINDS Human Cell and Data Repository (NHCDR). Fibroblasts were cultured in MEM medium supplemented with 15% fetal bovine serum.

### Cell culture, transfections and coimmunoprecipitation

SH-SY5Y cells were cultured in DMEM medium supplemented with 10% fetal bovine serum. Transient transfection with MYC-LRRK2 and HA-AP2A2/B1/M1/S1 or HA-AP2M1 truncated fragments, or HA-AP2M1 with Flag-LRRK2 truncated fragments was carried out using LipoD293 (Signagen) as per the manufacturer’s introductions. After 48 hours, cells were washed by PBS once, lysed in immunoprecipitation (IP) buffer (1% Triton X-100, 0.5% NP40, 150 mM NaCl, 20 mM HEPES, pH=7.4, 1 mM EGTA, 1 x Complete mini protease inhibitor cocktail (Pierce)) by rotation at 4 °C for 1 hour. Cell lysates were centrifuged at 15,000 rpm for 15 mins. Supernatants were incubated with protein-G Dyna beads (Bio-Rad) pre-coated with anti-MYC or anti-HA antibodies following by rotating overnight at 4 °C. The Dyna beads were pelleted and stringently washed five times with IP buffer supplemented with 500 mM NaCl. The immunoprecipitated proteins were resolved on SDS/PAGE and subjected to immunoblotting.

### *In vitro* Kinase assay

An *in vitro* kinase assay was performed as previously described (15, 25). GST tagged AP2M1-WT, T156A were expressed in *E. coli* BL21 and purified by Glutathione beads as per the manufacturer’s instructions (GE healthcare). The purity of the GST tagged proteins was assessed with denaturing SDS-PAGE followed by Coomassie blue staining. GST-LRRK2-WT, GS or DA recombinant proteins (970-2527aa) (Thermofisher), GST-AAK1 (1-501aa) (Thermofisher) with or without GST-AP2M1-WT or T156A protein were subjected to a kinase reaction in kinase buffer (20 mM HEPES pH=7.4, 150 mM NaCl, 10 mM EGTA pH=8.0, 20 mM β-Glycerol phosphate, 10 μM ATP, 0.5 μCi γ- ^32^P-ATP, 20 mM MgCl_2_). The reactions were incubated at 30 °C for 20 mins, put on ice, resolved on SDS-PAGE. LRRK2 autophosphorylation and AP2M1 phosphorylation were imaged using a Typhoon Phosphoimager. Input levels of protein were determined by immunoblotting of a GST-HRP antibody. The reactions without radioisotope were subjected to mass spectrometric analysis to identify AP2M1 phosphorylation sites by LRRK2.

### Mass spectrometric analysis

Mass spectrometry analysis was performed by the Taplin Biological Mass Spectrometry Facility (Harvard Medical School, MA, USA).

### Generation of LRRK2 knockout (KO) SH-SY5Y cell line by the CRISPR/Cas9 system

The gRNA (GAGTCCAAGACGATCAACAG) that target exon 2 of human LRRK2 genomic sequence was subcloned into plasmid pSpCas9(BB)-2A-Puro(PX459) V2.0 (a gift from Feng Zhang, Addgene, Plasmid #62988, http://n2t.net/addgene:62988; RRID:Addgene_62988) following previous protocol^(49)^. SH-SY5Y cells were transfected with LRRK2 gRNA plasmids. After 48 hours, single cell was plated and maintained in a 96 well plate. Deletion of LRRK2 was verified by Western blot with LRRK2 specific antibody and sequencing confirmation (Supplemental Figure 3). Multiple LRRK2 KO SH-SY5Y cell lines were selected.

### Subcellular fractionation

Cells were collected in fractionation buffer (20 mM HEPES pH=7.4, 10 mM KCl, 2 mM MgCl2, 1 mM EDTA, 1 mM EGTA, 1 x Complete mini protease inhibitor cocktail (Pierce)), incubated on ice for 15 mins following by passing through a 27-gauge needle 10 times and leaving on ice for 20 mins. The lysates were centrifuged at 8000g for 5 min. The resulting supernatant, which are the cytoplasm and membrane fractions, was further centrifuged in an ultracentrifuge at 100,000 g for 2 hours to produce the pellet (membrane fraction) and the supernatant (cytosol fraction).

### Liquid nitrogen coverslip freeze-thaw protocol

Liquid nitrogen coverslip freeze-thaw protocol to deplete cytosol was performed as previously described (18). Briefly, cells were plated on poly-L-ornithine coated coverslips, transfected with indicated DsRed-LRRK2 plasmids and transduced with lentiviruses carrying eGFP-AP2M1 WT or T156A. After 48 hours, cells were chilled on ice, washed twice with ice-cold PBS, and incubated in ice-cold glutamate buffer (25 mM KCl, 25 mM HEPES, pH=7.4, 2.5 mM magnesium acetate, 5 mM EGTA, 150 mM potassium glutamate). The coverslip was then dipped in liquid nitrogen for 5 s and allowed to thaw for few seconds, followed by gentle washes with ice-cold glutamate buffer and rehydration for 5 mins in ice-cold PBS on-ice. Cells were then fixed in cold 4% paraformaldehyde (PFA) for 20 mins, washed and mounted onto slides using Vectashield mounting medium (Vector Laboratories). Imaging was conducted on a Zeiss confocal 880 microscope (Carl Zeiss). The intensity of membrane-associated fluorescence was quantified by ImageJ software (NIH).

### Transferrin (Tfn) uptake Assay

Transferrin uptake assay was performed as previously described (50). Briefly, mouse primary hippocampal neurons from postnatal (1-day old) pups were plated on poly-L-ornithine coated coverslips in a 12 well plate and maintained in Neurobasal-A medium supplemented with B27 supplement and L-glutamine. After cultured for 5 days, cells were transduced with the indicated lentiviruses for 48 or 72 hours. The cells were then washed once with Neurobasal-A medium and incubated for 1 hour at 37 °C in Neurobasal-A medium without serum before incubated with 20 ug/mL Alexa Fluor 594–conjugated transferrin (Molecular Probes) for 16 mins at 37 °C. The cells were chilled on ice, washed once with ice-cold PBS, incubated with ice-cold acid wash buffer (0.2 M acetic acid, 0.5 M NaCl) for 5 mins on ice, then washed once with PBS and fixed with 4% PFA. Imaging was conducted on a Zeiss confocal 880 microscope (Carl Zeiss). The intensity of internalized Tfn-594 fluorescence in neuronal cell body was quantified by ImageJ software.

### Isolation of CCVs (clathrin-coated vesicles) and Uncoating Assays

CCVs were isolated as previously described(51, 52). Briefly, the homogenate from 10 whole mouse brains were centrifuged at 19,000g for 40mins. The resulting supernatant was centrifuged at 43,000g for 60 mins. The pellet containing the CCVs was resuspended in an equal volume of 12.5% Ficoll and 12.5% sucrose and centrifuged at 43,000g for 40min. The supernatant was then pelleted by centrifugation at 88,000 *g* for 1 hour at 4 °C. The pellet was resuspended in 4 ml homogenization buffer (100 mM MES, pH 6.5, 0.5 mM MgCl_2_, 1 mM EGTA, 1 mM DTT, and 0.1 mM PMSF) and layered over 6 ml of 8 % sucrose-D_2_O solution in the homogenization buffer and centrifuged at 100,000 *g* for 2 hours at 4 °C. The resulting pellet containing CCVs was collected, washed, and resuspended in the homogenization buffer. The suspension was further centrifuged at 20,000 *g* for 10 mins at 4 °C and the resulting supernatant containing CCVs was collected, snap frozen in aliquots and stored at −80 °C for uncoating assays.

Uncoating assays were performed as previously described(21, 51). 5–6 μg of above purified CCVs were incubated with recombinant proteins GST-LRRK2 G2019S or D1994A, GST proteins as a control, in an ATP-regenerating system (1.3 μg Hsc70, 0.8 mM ATP, 5 mM of creatine phosphate, and 0.2 IU of creatine phosphokinase, 10 mM ammonium sulfate, 20 mM HEPES, pH 7.0, 2 mM magnesium acetate, and 25 mM KCl) in a final volume of 50 μl. The mixture was incubated for 10 mins at 25 °C, put on ice, centrifuged at 100,000 *g* for 20 mins at 4 °C. Pellet were resolved by SDS-PAGE and coated AP2M1 on CCVs was analyzed by Western blot.

### Preparations of Lentiviral AP2M1-WT (or TA)-eGFP

The second-generation lentiviral packaging system was employed to produce high-titer lentiviruses. Briefly, pFUGW-AP2M1-WT-eGFP and pFUGW-AP2M1-T156A-eGFP lentiviral plasmids were transfected into HEK 293FT cells along with viral packaging plasmids (psPAX2 and pMD2.G). After 48 hours, the culture media were collected, and viral particles were precipitated by centrifugation at 35,000 g for 2 hours. Viral particles were resuspended into serum free medium and stored at −80 °C.

### Primary neuronal cultures and viability assay

Primary cortical neuronal cultures were prepared from embryonic day 15–16 CD1 fetal mice (Charles River) (14, 25). Briefly, Cortices were dissected and dissociated by Trypsin (Invitrogen). The cells were seeded into 24-well plates pre-coated with poly-L-ornithine and were maintained in Neurobasal medium (Invitrogen) supplemented with B27 supplement and L-glutamine. The glial cells were inhibited by adding 5-fluoro-20-deoxyuridine (5F2DU, 30 μM, Sigma) at days *in vitro* (DIV) 4. To assess LRRK2 and AP2M1-induced toxicity, MYC-LRRK2, HA-AP2M1 and eGFP were co-transfected at a plasmid ratio of 10:10:1 into neurons at DIV 5 using Lipofectamine 2000 reagent (Invitrogen). At 48 hours post-transfection, neurons were fixed with 4% PFA. Fixed cells are washed with PBS, permeabilized and blocked for 1 hour with PBS containing 5% goat serum and 0.3% Triton X-100, then detected with rabbit anti-GFP antibody followed by AlexaFluor-488 anti-rabbit IgG antibody. The stained cells were further labelled by TUNEL (TdT-mediated dUTP-X nick end labeling) with an In-Situ Cell Detection Kit (TMR red) from Roche (12156792910). Fluorescent images were collected on a Zeiss Automatic stage microscope with Zen blue software. Neurons with no obvious neurite process and TUNEL staining positive were counted as non-viable cells. For each independent experiment, the percent viability of GFP-positive neurons (*n* >100) was determined and normalized to the control neurons transfected with pEGFP-N1 plus pcDNA3.1 empty vector.

### Immunocytochemistry

Neurons transfected with indicated plasmids were fixed in 4% PFA for 20 mins, washed with PBS, permeabilized and blocked for 1 hour with PBS containing 5% goat serum and 0.3% Triton X-100. The neurons were then incubated with mouse anti-MAP2 antibody at 4 °C overnight. After washing with PBS, the neurons were incubated with goat anti-mouse IgG conjugated to CY5 at room temperature for 2 hours. The coverslips were mounted onto slides using Vectashield mounting medium (Vector Laboratories). The imaging was conducted on a Zeiss confocal 880 microscope with Zen black software or a Zeiss Automatic stage microscope with Zen blue software.

### TIRF (total internal reflection fluorescence) microscopy

TIRF imaging experiments were performed on an inverted wide-field microscope operated in the TIRF mode as described previously in detail(53). Briefly, cells were plated on poly-L-ornithine coated coverslips, transfected with indicated DsRed-LRRK2 and eGFP-AP2M1 plasmids. After 48 hours, cells were fixed in 4% PFA for 20 mins, washed and mounted onto slides using Vectashield mounting medium (Vector Laboratories). Imaging was conducted on a TIRF microscope.

### *Drosophila* genetics

*Ddc-GAL4* and *UAS-GFP* fly lines were obtained from Bloomington Stock Center. pUAST-HA-AP2M1, T156A, and pUAST-attB-LRRK2 WT and G2019S were microinjected into *Drosophila* embryos (BestGene Inc.). The transgenic AP2M1 were crossed with flies carrying LRRK2 different forms. The resulting bigenic flies were crossed with *Ddc-GAL4>UAS-GFP*, which induces the co-expression of GFP, AP2M1 and LRRK2 in dopamine and serotonin neurons.

### Statistical analysis

Statistical analysis was performed with Prism 6.0 software (GraphPad). One-way ANOVA followed by a Tukey’s post hoc test was used for data analysis for multiple comparisons and Student’s *t* tests (unpaired, two-tailed) was used for two comparisons as described in the figure legends. Data represent mean ± SEM, and *p* ≤ 0.05 was considered statistically significant. \**p* <0.05, \*\**p* < 0.01, \*\*\**p* < 0.001, \*\*\*\**p* < 0.001.

### Study Approval

All procedures involving animals were approved by and conformed to the guidelines of the Institutional Animal Care Committee of Kansas State University.

## Author Contributions

Q.L. performed the experiments; Q.L., J.Y. and Y.X. analyzed Data; Y.X. and J.Y. conceived the study, designed the experiments and wrote the manuscript, J. B. G. and D.H. provided assistance on the TIRF microscopy study.

## Acknowledgements

This work was supported, in part, by grants from NIH/NINDS R01 NS112506, NIH/NIA K01 AG046366 award, Parkinson’s Foundation Stanley Fahn Junior Faculty award PF-JFA-1934, American Parkinson Disease Association (APDA) Research Grant, NIH/NIGMS P20 GM113109 pilot grant, SUCCESS-FYI Intramural Grant from Kansas State University College of Veterinary Medicine. The authors acknowledge Confocal Core and Molecular Biological Core supported by CVM-KSU. The authors thank Dr. Ping Li of the Department of Chemistry (KSU) for assistance on MS data analysis.

## Notes

The Authors have declared that no conflict of interest exists.

### Competing Interest Statement

The authors have declared no competing interest.

### Summary of Updates

The abstract revised.

