## Supplementary material for "LRRK2 regulates AP2M1 phosphorylation cycles to mediate endocytosis and dopaminergic neurodegeneration"

Supplemental Fig. 1

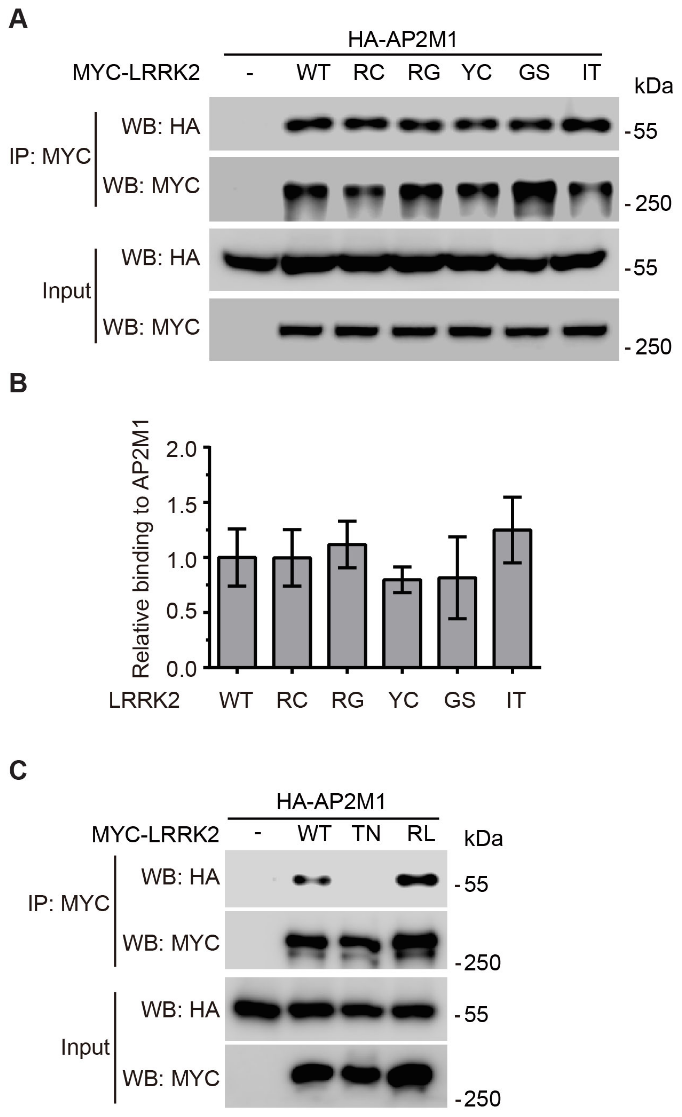


**Supplemental Figure 1. AP2M1 interacts with LRRK2 mutants. A.** Cell lysates from HEK 293T cells co-transfected with HA-AP2M1 and WT or familial mutants MYC-LRRK2 (R1441C/G, Y1699C, G2019S, I2020T) were subjected to Co-IP with anti-MYC followed by anti-HA immunoblotting or by anti-MYC to show an equivalent amount of input. **B.** Quantification of different mutant LRRK2 binding to AP2M1 relative to WT-LRRK2. Bars represent the mean ± SEM (n=3). Statistical significance was determined by One-way ANOVA followed by a Tukey's post hoc test. No significant difference was found between WT and mutant LRRK2 in terms of binding to AP2M1. **C**. Cell lysates from HEK 293T cells co-transfected with HA-AP2M1 and WT or GTP binding mutants MYC-LRRK2-T1348N (TN) and MYC-LRRK2-R1398L (RL) were subjected to Co-IP with anti-MYC followed by anti-HA immunoblotting or by anti-MYC to show an equivalent amount of input.

Supplemental Fig. 2

**A**

migglfiynh kgevlisrvy rddigrnavd afrvnvihar qqvrspvtni
artsffhvkr sniwlaavtk qnvnaamvfe flykmcdvma ayfgkiseen
iknnfvliye lldeildfgy pqnsetgalk tfitqqgiks qhqtkeeqsq
itsqv**T**gqig wrregikyrr nelfldvles vnllmspqgq vlsahvsgrv
vmksylsgmp eckfgmndki viekqgkgta detsksgkqs iaiddctfhq
cvrlskfdse rsisfippdg efelmryrtt kdiilpfrvi plvrevgrtk
levkvviksn fkpsllaqki evriptplnt sgvqvicmkg kakykasena
ivwkikrmag mkesqisaei ellptndkkk warppismnf evpfapsglk
vrylkvfepk lnysdhdvik wvryigrsgi yetrc

**B**

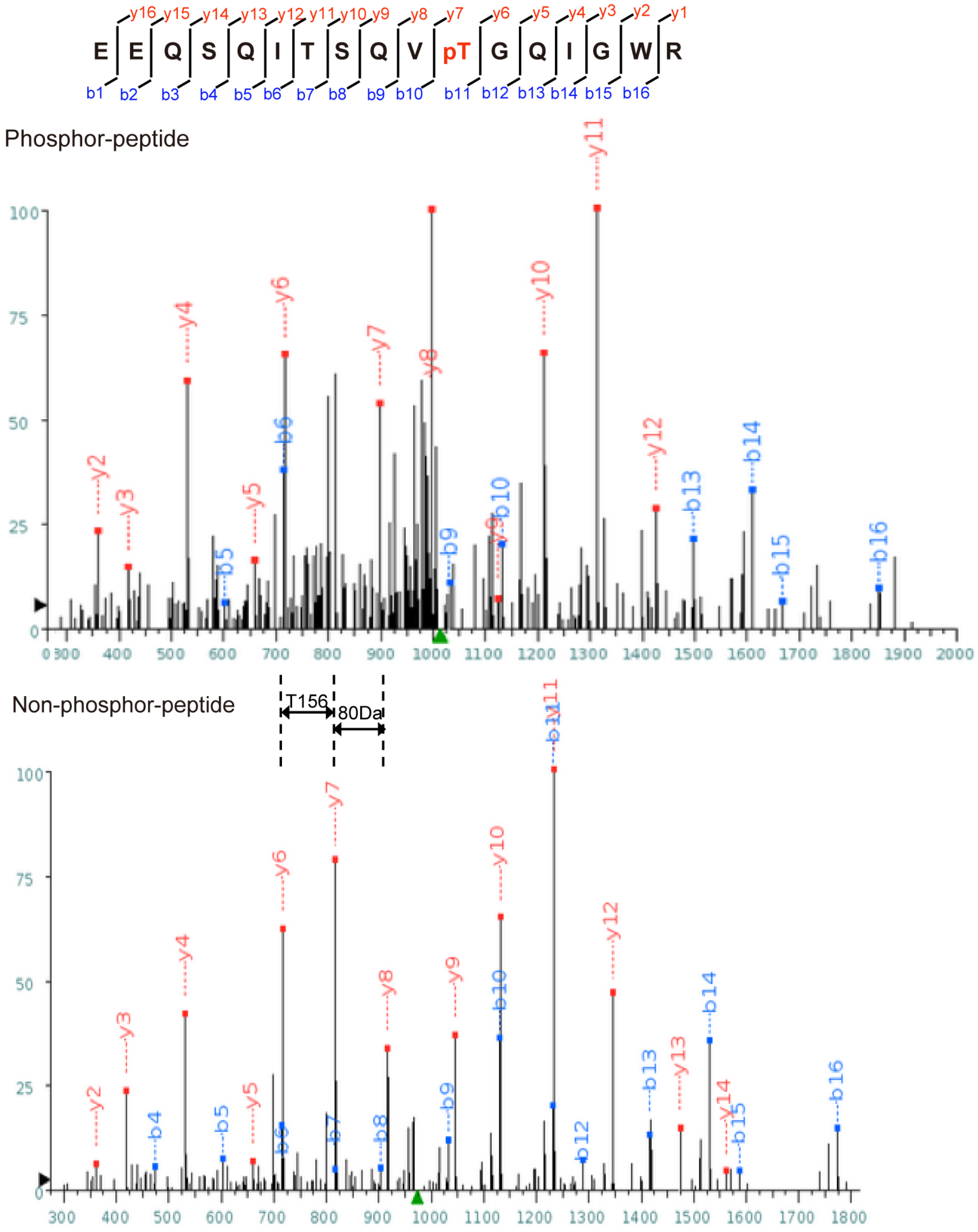

**Supplemental Figure 2. Mass spectrometric analysis of AP2M1 phosphorylation sites by LRRK2. A.** AP2M1 is phosphorylated by LRRK2 majorly at threonine T156. Mass spectrometric analysis reveals 95.3% sequence coverage of AP2M1. Phosphorylated sites are indicated in red. **B.** Both nonphosphorylated and phosphorylated AP2M1 were subjected to LC-MS/MS to identify the phosphorylation site (bottom panel). LC-MS/MS spectra of the nonphosphorylated peptide (EEQSQITSQVTGQIGWR) and the phosphorylated peptide (EEQSQITSQVpTGQIGWR) are compared, demonstrating that there is the 80-Da shift for the T156 ion containing the phosphate moiety (y7). The phosphorylated amino acid is preceded by a "p" and is highlighted in red.

Supplemental Fig. 3

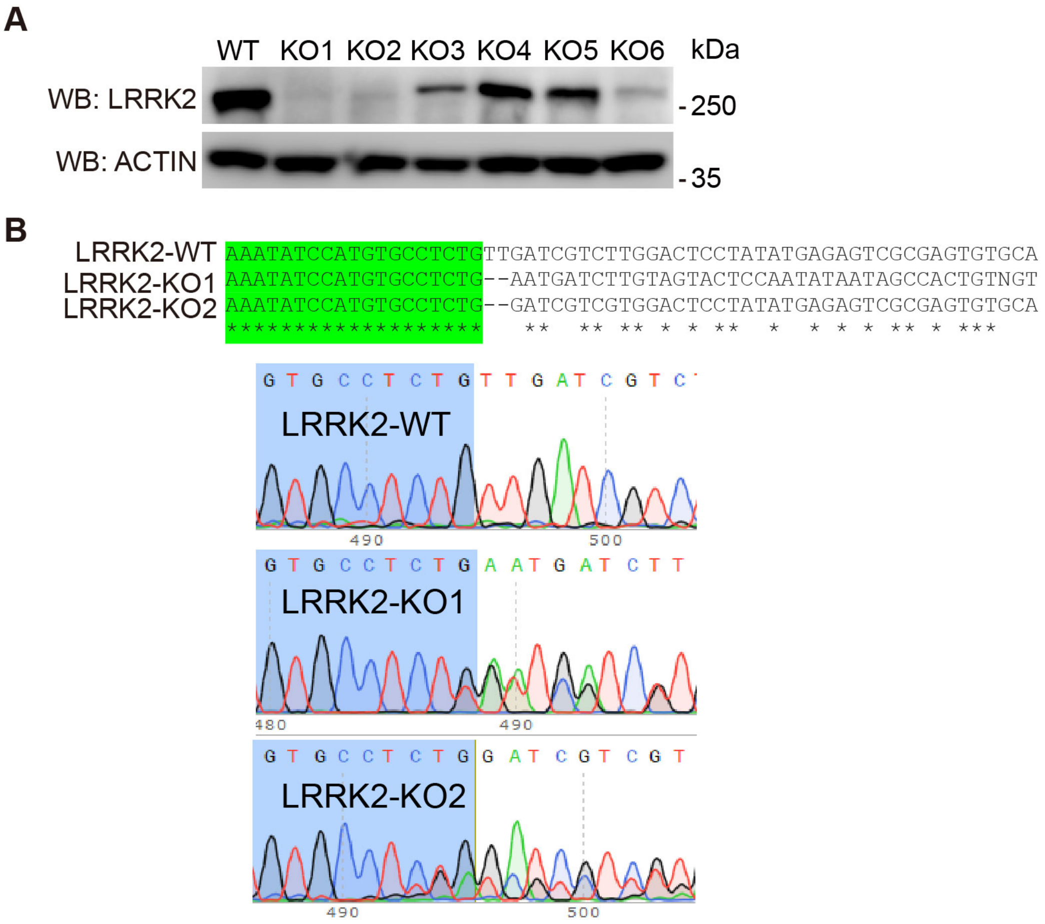


**Supplemental Figure 3. Generation of LRRK2 knockout (KO) SH-SY5Y cells by the CRISPR/Cas9 system. A.** LRRK2 protein level was completely depleted in LRRK2 KO cells. Western blot analysis of endogenous LRRK2 levels by immunoblotting with anti-LRRK2. **B.** Sequencing of the target region confirmed a frame shift of LRRK2 encoding sequences.

Supplemental Fig. 4

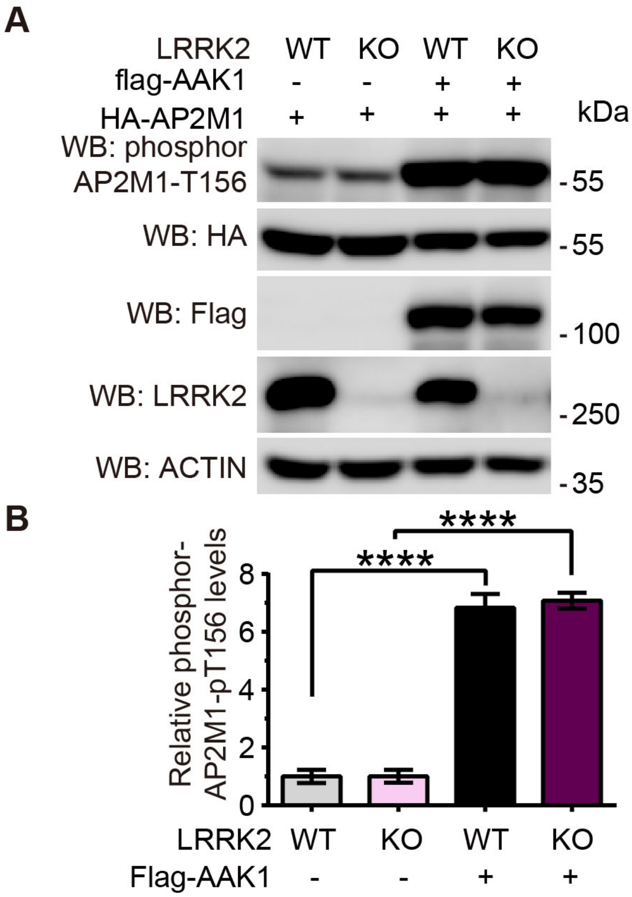


**Supplemental Figure 4. AAK1 phosphorylates AP2M1 independently of LRRK2. A.** AAK1 and AP2M1 were overexpressed in LRRK2 WT and KO cells. Lysates were subjected to immunoblotting with anti-pT156-AP2M1. **B.** Quantification of overexpressed AP2M1 phosphorylation level in LRRK2 WT and KO cells with overexpression of AAK1. Statistical significance was determined by Student's *t* tests. Bar represents the mean ± SEM, n=5, ****$P < 0.0001$.

Supplemental Fig. 5

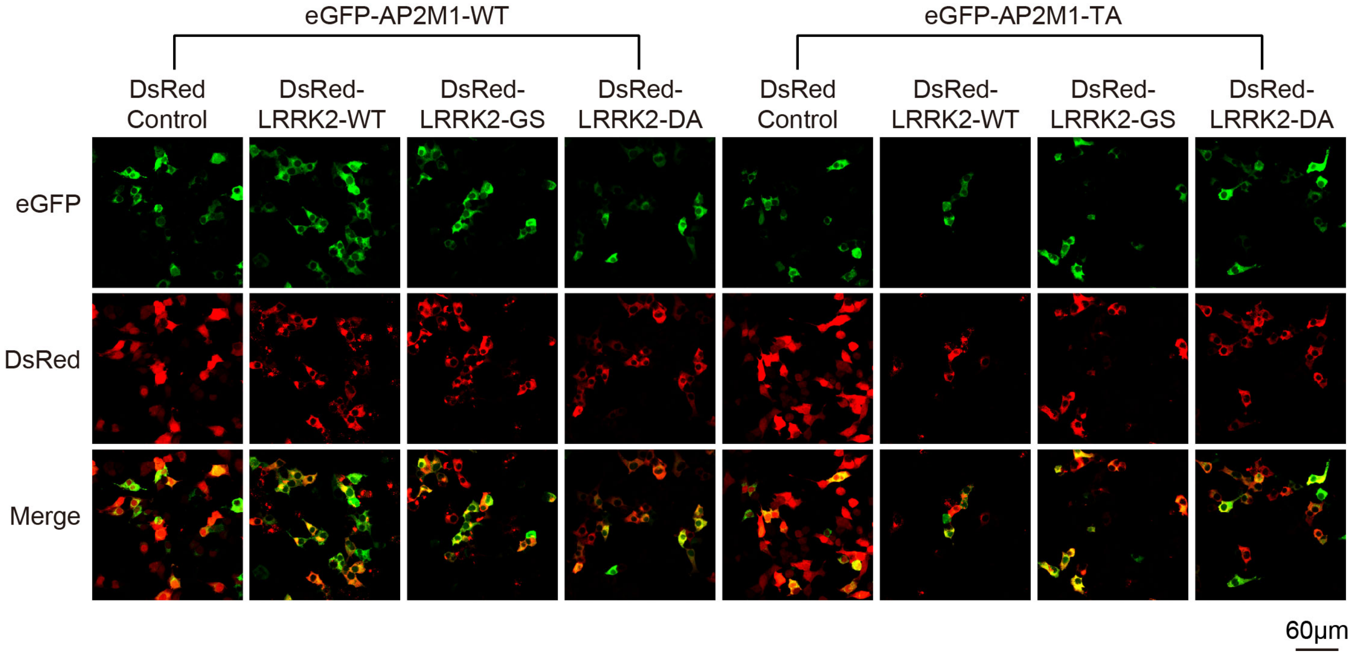


**Supplemental Figure 5. Co-transfection of AP2M1-eGFP and DsRed-LRRK2 in SH-SY5Y cells at a plasmid ration of 1:5.** >90% of eGFP (AP2M1) positive cells were DsRed (LRRK2) in TIRF microscopy study (Figure 4d) as verified by confocal microscopy.

Supplemental Fig.6

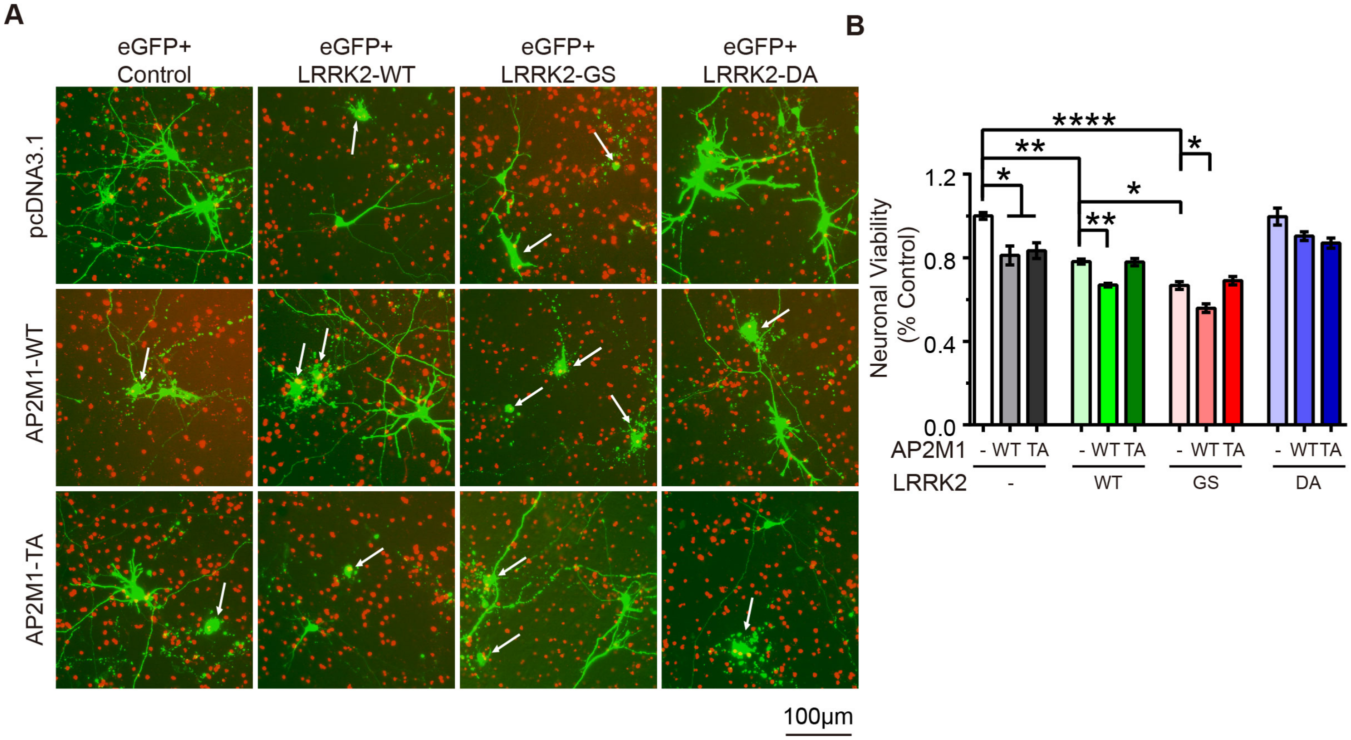


**Supplemental Figure 6. AP2M1 WT enhances LRRK2 induced neuronal toxicity. A.** Representative fluorescent images (green) showing that mouse primary cortical neurons were co-transfected with MYC-LRRK2, HA-AP2M1 with eGFP at a plasmid ratio of 10:10:1 at DIV 5. Neuronal viability was analyzed at 48-hour post-transfection with non-viable neurons exhibiting no obvious neurite process and positive TUNEL staining (arrow). Scale = 100 μm. **B.** Quantification of neuronal viability. Bars indicate the viability (n > 100) for each transfection condition expressed as a percent of control neurons (eGFP with pcDNA3.1 empty vector). Statistical significance was determined by one-way ANOVA followed by a Tukey's post hoc test. Data represent the mean ± SEM from three independent experiments with n > 100 cells quantified in each experiment. **P* < 0.05, ***P* < 0.01, *****P* < 0.0001.
